# Interactions with lipid membrane modulate the conformational dynamics and energy landscape of Tumor Necrosis Factor

**DOI:** 10.1101/2025.11.28.690881

**Authors:** George Khelashvili, István Kolossváry, Woody Sherman, Fabio Trovato

## Abstract

Tumor Necrosis Factor (TNF) is a trimeric cytokine that exists in soluble (sTNF) and membrane-bound (mTNF) forms, both of which regulate immune responses through interactions with their cognate receptors. sTNF has been the subject of extensive biophysical investigations, leading to a prevailing mechanistic model in which a symmetric arrangement of the trimer promotes receptor signaling, whereas asymmetric conformations inhibit this function. In contrast, the structural and energetic landscape of mTNF remains largely under-explored. Here, we combined multi-microsecond unbiased molecular dynamics and Metadynamics-Adaptive Biasing Force simulations to characterize mTNF embedded in a heterogeneous lipid bilayer. We show that in the absence of membrane engagement, the extracellular domain (ECD) dynamically samples multiple asymmetric conformations similar to those observed in sTNF. The association of the ECD with the membrane, mediated primarily by the basic residues R78–R82, R107, R108, R120, and R207, restricts conformational heterogeneity and stabilizes the symmetric state. By quantifying the ECD– membrane interactions, we demonstrate that the symmetric state in the membrane-bound ECD is stabilized over asymmetric conformations to a greater extent than in sTNF. This energetic effect, together with the spatial confinement imposed by the lipid bilayer, may explain the previously observed reduced affinity of certain biologics for mTNF. In conclusion, our work elucidates how the membrane influences the structure, dynamics, and energetics of TNF. These mechanistic insights could guide future efforts to design mTNF-selective inhibitors that account for both membrane constraints and the energetic modulation of the ECD conformational landscape.

**STATEMENT OF SIGNIFICANCE:** Tumor necrosis factor (TNF) drives inflammation and is one of the most successful drug targets in medicine. Most of what is known concerns the soluble form; the membrane-anchored form, which signals between cells and is less susceptible to several approved biologics, has remained largely uncharacterized. Using multi-microsecond atomistic and enhanced-sampling simulations of TNF embedded in a lipid bilayer, we show that its extracellular domain binds the membrane through a defined set of basic residues and that this interaction suppresses the asymmetric, receptor-incompetent conformations that the soluble form samples. Therefore, our study shows how the membrane modulates TNF conformational landscape. These computational results offer a mechanistic explanation for the reduced potency of some TNF biologics and may support future efforts in designing form-selective inhibitors.

## INTRODUCTION

TNF is a pro-inflammatory cytokine that belongs to the Tumor Necrosis Factor superfamily and is expressed in several cell types as a transmembrane type II polypeptide (mTNF)^1^. Upon enzymatic cleavage, the extracellular domain (ECD) is released as a soluble form (sTNF)^2^. Binding of sTNF to receptor TNFR1 can activate the inflammation, proliferation or apoptosis pathways, depending on conditions such as TNF oligomerization and cluster formation^3–5^. Accumulating evidence suggests that not only sTNF, but also mTNF participates in inflammatory processes by interacting with receptors TNFR1 and TNFR2^1,6^

At physiological concentrations, sTNF is a trimer composed of units with an elongated, “jelly-roll” topology^7^. As demonstrated by double electron–electron resonance (DEER) experiments^8^, molecular dynamics (MD) simulations^9^, and a conformation-selective monoclonal antibody^10^, sTNF is in equilibrium between different trimeric conformational states. In the first state, the three monomers assemble symmetrically, creating solvent-exposed grooves between the inter-subunit interfaces^7^, each capable of binding one receptor monomer with high affinity^5^. The second state is asymmetric and is characterized by a partial widening of one of the grooves^10,11^, which reduces receptor binding and therefore causes inhibition of sTNF function^12^. TNF groove widening has been observed primarily in two contexts: co-crystal structures containing small molecules bound at the center of the trimer, which effectively lock sTNF in a receptor-incompetent state; DEER-informed models that capture transient excursions of apo TNF into asymmetric states. Two other members of the TNF superfamily, CD40L^13^ and TRAIL^14^, have been shown to populate non-symmetric states in different conditions.

Despite the extensive knowledge on sTNF, much less is known about mTNF, limiting our mechanistic understanding of the biophysical and clinical behaviors of the two forms. For example, some biologics and small molecule inhibitors bind mTNF but show limited or no inhibition^15,16^. It is also unclear why the most widely used biologics bind mTNF more weakly than to sTNF^17^, or why antibodies neutralizing sTNF cannot inhibit liposome-associated TNF^18^. Previous experimental and computational studies investigated the interactions between mTNF and TNFR1 and their consequences on receptor clustering ^4,19^. However, these studies did not investigate the mTNF conformational equilibria relevant to the inhibition of the TNF-receptor interactions. These knowledge gaps motivated us to study whether and how the plasma membrane can modulate structure and dynamics of TNF, therefore providing a mechanistic interpretation to the experimental observations.

In this work, we sought to characterize the conformational dynamics and energetic landscape of mTNF in a heterogeneous lipid bilayer ^20^, and compare them to sTNF. To this end, we integrated molecular modeling, multi microsecond unbiased molecular dynamics simulations and Metadynamics coupled with extended Lagrangian Adaptive Biasing Force (Meta-eABF) simulations^21^. We first validated our approach on sTNF by benchmarking against available experimental and computational data, before applying the same framework to mTNF.

Our study reveals a heterogeneous ensemble of low-energy symmetric and asymmetric conformations explored by sTNF, in agreement with previous reports^8,16^, whereas the mTNF free energy landscape is dominated by an ensemble of symmetric conformations in our simulations of mTNF engaged with the membrane. Simulations further predict specific modes of interaction between the mTNF ECD and the membrane. In a long-lived state, the ECD associates stably with lipids through residues R78-R82, R107, R108, R120, and R207, which are known to participate in TNF receptor recognition^6,22^. Importantly, during this membrane-bound configuration, the trimer remains in the symmetric conformation. These findings demonstrate that the mTNF ECD can directly engage the plasma membrane, consistent with experimental observations for other TNF superfamily members, such as TRAIL and GITRL^14,23^.

In conclusion, we provide a detailed structural and energetic characterization of the mTNF conformational landscape, highlighting the specific mechanisms of ECD stabilization mediated by the lipid bilayer as well as differences with the sTNF conformational landscape. Our work may offer new insights for designing inhibitors that selectively target the two TNF forms.

## METHODS

Our computational pipeline (**Fig. 1**) consists of 4 major stages: i) generation of atomistic models for the symmetric and asymmetric mTNF trimer; ii) unbiased ensemble MD simulations of these models in atomistic lipid membranes to sample protein-membrane interactions and intra-protein conformational transitions; iii) analysis of the MD data using dimensionality reduction to select conformations of the system for subsequent biased sampling MD; and iv) performing enhanced sampling simulations to predict free energy surfaces. The methodological aspects of these 4 steps are described in detail below.

**Figure 1:**
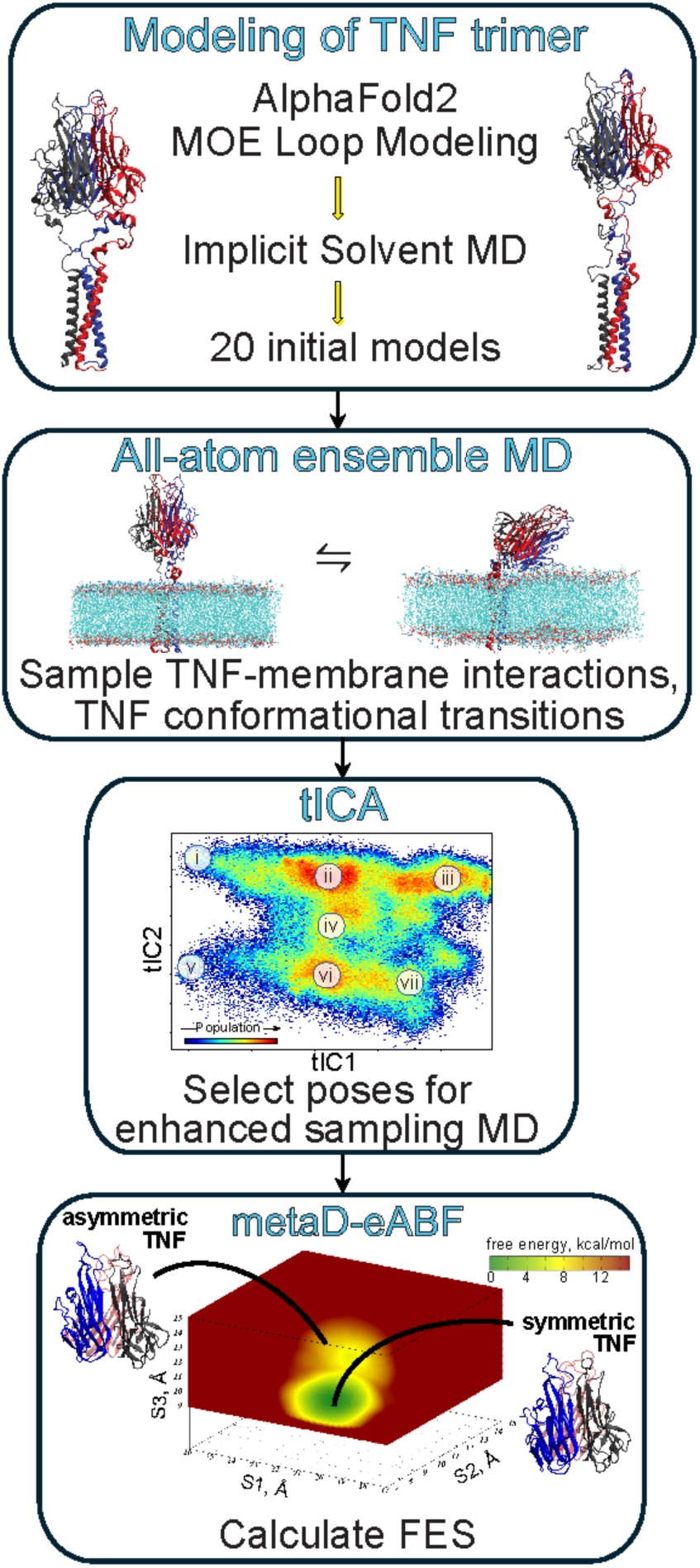
Multi-stage computational pipeline for studying mTNF conformational transitions and membrane interactions. The major stages of the pipeline are: i) generation of structural models for symmetric and asymmetric mTNF using a combination of protein structure prediction and implicit solvent MD simulation approaches; ii) unbiased ensemble MD simulations of the protein models in atomistic lipid membranes to sample protein-membrane interactions and intra-protein conformational transitions; iii) dimensionality reduction analysis of the MD data with tICA to select conformations of the system for subsequent biased sampling MD; and iv) biased Meta-eABF simulations to predict free energy surfaces. For all the methodological aspects of these 4 steps see Methods.

### Generation of structural models for mTNF

In this section we describe the three steps we used to model the mTNF conformations. In step 1, we model the TNF trimer based on the available X-ray structures; in step 2, we increase model diversity; in step 3, we embed the modeled systems in a heterogeneous lipid bilayer.

#### 1. Modeling of the TNF trimer

A single polypeptide of the TNF trimer is composed of 233 residues (Uniprot ID P01375). Proteolytic cleavage between residues 76 and 77 of the TNF polypeptide separates an extracellular soluble domain (sTNF) comprising residues 77 to 233 from the transmembrane (TM) domain (**Fig. 2a**). While the full-length TNF structure has not been experimentally determined, the structures of the sTNF (both in symmetric and asymmetric states) as well as of a monomeric TM helix have been resolved. To model symmetric and asymmetric trimer models of sTNF, we used crystal structures with the PDB IDs 1TNF and 7JRA, respectively^7,24^. In the latter, the asymmetric trimer is in complex with an inhibitor which was removed for our studies.

**Figure 2:**
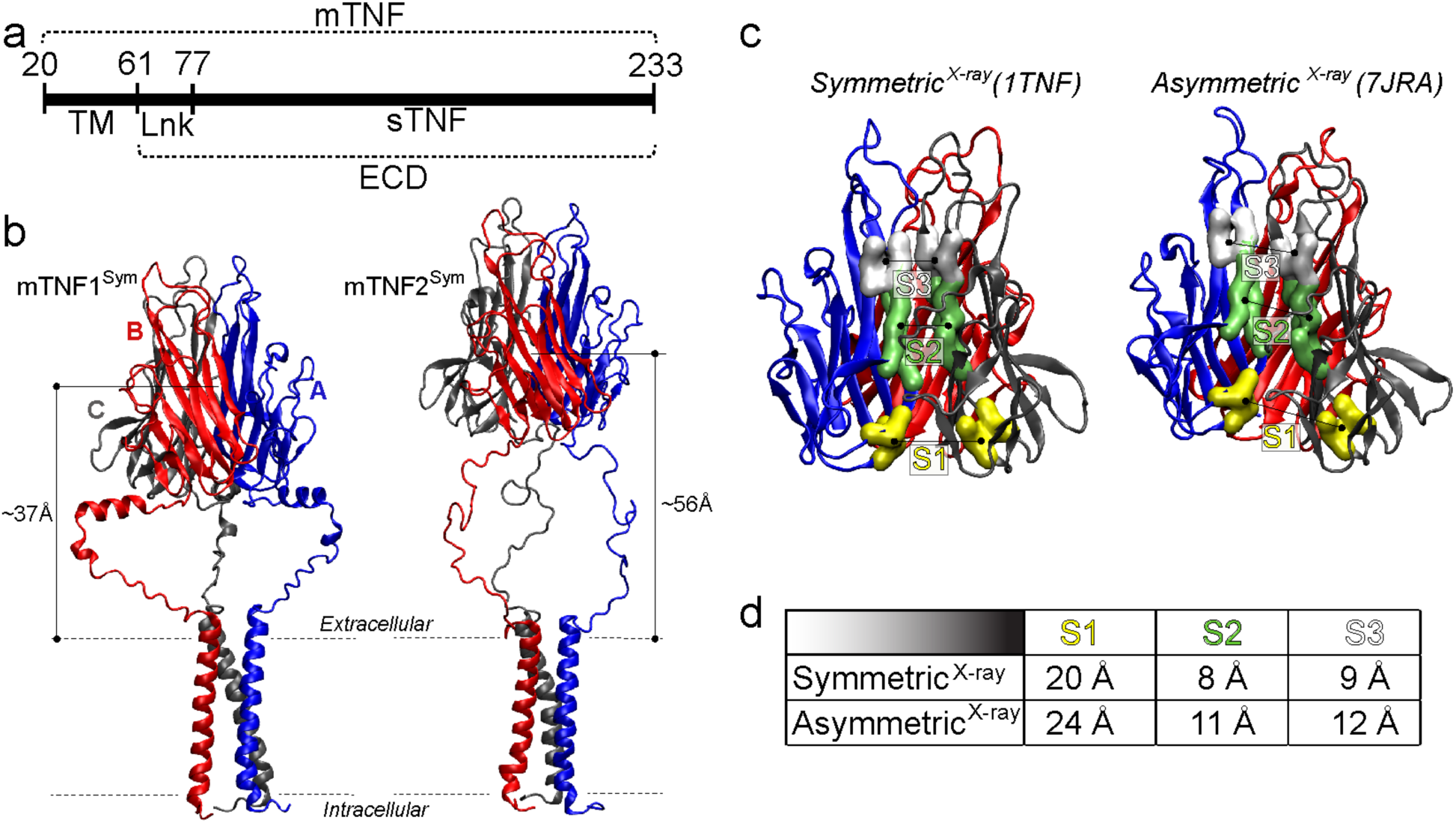
Structural models of mTNF. **(a)** Full TNF sequence highlighting the transmembrane (TM), linker (Lnk) and ECD regions. (**b**) Two structural models of symmetric mTNF, differing in the position of their ECD with respect to the membrane surface. The three subunits (A, B, and C) of the protein are shown in blue, red, and grey cartoons, respectively. Approximate membrane boundaries are marked as dotted lines. **(c)** Structures of experimentally determined models of symmetric (PDBID 1TNF) and asymmetric (PDBID 7JRA) sTNF. The color code is the same as in panel b. The yellow, green, and white surfaces represent the backbone atoms of the residues used for the definition of the S1, S2, and S3 distances used in the simulations (see Methods). **(d)** The values of the S1, S2, and S3 distances calculated from the two crystallographic structures shown in panel c.

The NMR structure deposited under PDBID 7ASY^25^ comprises a single helix of the TM domain. We used its amino acid sequence, corresponding to TNF residues 28 to 60, to construct a trimeric TM model. Specifically, we (i) used the sequence of residues 28 to 60 as input to Alphafold2-Multimer^26^, (ii) minimized the resulting model using Molecular Operating Environment (MOE)^27^, and (iii) predicted its free energy of association using the TMPfold webserver (https://opm.phar.umich.edu/tmpfold_server)^28^. As shown by the high pLDDT score (**Fig. S1a)**, Alphafold2-Multimer predicted a trimeric helix bundle with high confidence. The calculated free energy of association of -14.2 kcal/mol further supports the stability of the predicted trimeric TM domain. This prediction is consistent with size exclusion chromatographic experiments showing that the TNF TM region supports trimeric assembly under the tested experimental conditions^29^.

The positioning of the predicted TM domain within a model membrane was obtained using the OPM webserver (opm.phar.umich.edu)^30^. To extend the N-termini farther from the membrane, while keeping the simulation box at a reasonable size (see below), we decided to add residues 20 to 27 to each helix, in a disordered conformation, while omitting disordered residues 1 to 19 (**Fig. S1c)**. Although segment 1-19 contains a sequence implicated in TNF reverse signaling^31^, its omission from the model is not expected to influence the results of our simulations because we study processes taking place at the extracellular side. ^31^

Next, we connected the TM domain with the sTNF structures (symmetric or asymmetric) through a linker region. To increase the diversity of the structures employed to start the simulations described below, we used two different methods.

In the first method, the structure of amino acid sequence 58 to 81 was predicted using AlphaFold2^32^. Subsequently, the predicted linker was overlayed onto the TM and sTNF domains. Residues 58, 59 and 60 were eliminated from the Alphafold2 models and the linker residue 61 was connected to residue 60 of the TM domain. Since residues 77 to 81 are present both in sTNF and the linker, we eliminated residues 77 to 81 in sTNF and connected the linker residue 81 to residue 82 of sTNF. The two final full-length models were termed mTNF1^Sym^ and mTNF1^Asym^, where “1” indicates the method and superscripts “Sym” and “Asym” denote the symmetric and asymmetric trimer conformations, respectively.

In the second approach, we started with the mTNF1^Sym^ and mTNF1^Asym^ models, eliminated residues 61 to 81, increased the vertical distance between the TM and sTNF domains by roughly 20Å, corresponding to the length of the fully unfolded linker, and rebuilt residues 61 to 81 in a disordered conformation, using MOE loop modeling functionality. The resulting symmetric and asymmetric models were termed mTNF2^Sym^ and mTNF2^Asym^ respectively.

The four models, mTNF1^Sym^, mTNF1^Asym^, mTNF2^Sym^ and mTNF2^Asym^ were minimized. The definitions of the TM, linker and sTNF domains are shown in **Fig. 2a**. The 3D full-length models mTNF1^Sym^ and mTNF2^Sym^ are depicted in **Fig. 2b**, and the 3D models mTNF1^Asym^ and mTNF2^Asym^ can be found in **Fig. S1b**, labeled “AlphaFold” or “MOE loop modeling”.

#### 2. Implicit solvent MD simulations to increase model diversity

Implicit solvent MD simulations were carried out to enable us to efficiently generate diverse starting conformations of the mTNF constructs at all-atom resolution. At this stage, the goal was to generate a set of conformations with plausible linker diversity and mTNF ECD positioning with respect to the membrane. As described below, subsequent multi-microsecond MD simulations, performed in explicit solvent, account for solvent-mediated effects.

The Implicit solvent MD simulations were carried out in the absence of membrane and were started from the four mTNF1^Sym^, mTNF1^Asym^, mTNF2^Sym^ and mTNF2^Asym^ models built as described section “*Modeling of the TNF trimer*” above. The implicit solvent simulation protocol described below uses the OpenMM 7.7 library.

The initial coordinates and parm7 files were obtained using AMBER LEaP, with (i) ff14SB force field and the generalized Born radii model mbondi3^33^. A heating phase was carried out by gradually heating the system from 0 to 310 K in 10K intervals, each interval run for 0.12 ns, followed by a production phase of 1 μs at 310K. We additionally used hydrogen mass repartitioning^34^, an integration timestep of 4 fs, Langevin thermostat with friction of 1.0 ps^-1^ and harmonic positional restraints with force constant of 100 kJ·mol^-1^·nm^-2^ on every backbone atom but the linker domain. For each of the four mTNF models, we ran 5 replicates.

To select representative full-length structures from the implicit solvent simulations, we used GROMACS clustering tool. We combined the 5 simulation replicates of each mTNF model into a single trajectory file with 25,000 frames. The linker segments (residues 61 to 81 in each TNF monomer) from the combined trajectory were then clustered using the Root Mean Square Deviation (RMSD) of linker Cα atoms and an RMSD cut-off value of 5 Å. 16 representative structures within the top four clusters were extracted. Thus, together with the initial AlphaFold- and MOE-based models, we created 20 mTNF models, as depicted in **Fig. S1b** (5 for mTNF1^Sym^, 5 for mTNF2^Sym^, 5 for mTNF1^Asym^, and 5 for mTNF2^Asym^).

#### 3. Construction of membrane-embedded mTNF systems

Using Membrane Builder in MolCube, a commercial version of CHARMM-GUI ^35^, each of the 20 mTNF models we built was inserted into a lipid membrane with sizes 120 Å x 120 Å (**Fig. S1b**). Because membrane TNF is expressed by a broad range of cell types, including monocytes/macrophages, lymphocytes, NK cells, dendritic cells, endothelial cells, and fibroblasts, its native lipid environment is expected to vary with cellular context. We therefore used a representative plasma-membrane asymmetric lipid composition^20^ as a physiologically motivating approximation. Thus, the model membrane contained a 48:45:3:4 mixture of SM:POPC:POPE:POPS lipids on the extracellular (EC) leaflet and a 3:25:42:30 mixture of the same lipids on the intracellular (IC) leaflet^20^. Phosphatidylinositol lipids, which are usually present in small (2%) concentration on the IC leaflet of plasma membranes, were not considered here because at the time of this work reliable AMBER Lipid21-compatible force field parameters for these lipids were not available. The absence of the negative charged of phosphatidylinositol lipids was compensated here by considering appropriately elevated concentration of POPS lipids on the IC leaflet. We also did not include cholesterol in our lipid mixture, as its presence would necessitate significantly longer simulation times to properly equilibrate the membranes. As we describe in the Results section, our main findings relate to the mTNF ECD engaging with the lipid membrane via positively charged residues. The presence of cholesterol would likely strengthen these interactions^36^ without affecting the mechanistic insights we describe.

The final 20 mTNF-membrane complexes were solvated and ionized with 150 mM KCl salt, resulting in system sizes ranging between ∼320,000 and ∼350,000 atoms.

### Molecular dynamics simulations

We performed two sets of MD simulations (**Fig. 1**): 1) multi-microsecond unbiased MD simulations to gain insights into the dynamics of mTNF and to select suitable frames as inputs for subsequent enhanced sampling MD; and 2) Enhanced sampling meta-eABF simulations to compute the free energy surfaces along pre-determined Collective Variables (CVs).

#### Unbiased MD Simulations

The 20 modeled mTNF systems (**Fig. S1b**) were first equilibrated using the protocol provided in MolCube, which mirrors the standard 6-step equilibration protocol used in CHARMM-GUI^35^. During this stage, the backbone of the protein was first harmonically constrained and subsequently gradually released in three steps of 5 ns each, changing the restrain force constants from 1, to 0.5, and 0.1 kcal/ (mol Å2), respectively. After this equilibration phase, each system was subjected to unbiased 1 μs-long MD simulations. The simulations were carried out with OpenMM version 7.7^37^ using AMBER ff14SB and Lipid21 force fields for the protein and lipids, respectively, and the TIP3P water model^33,38^. Hydrogen mass repartitioning was applied to the system^34^. The simulations implemented PME for electrostatic interactions and were performed at 310 K temperature, under NPT ensemble using semi-isotropic pressure coupling, and with 4fs integration time-step. Monte Carlo barostat and Langevin thermostat were used to maintain constant pressure and temperature, respectively. Additional parameters for these simulations included: friction coefficient of 1.0/picosecond, Ewald tolerance of 0.0005, and with “Hbonds” constraints. For the van der Waals interactions, we applied a cutoff distance of 12 Å, switching the potential from 10 Å.

##### CV definition and protocol for Meta-eABF simulations

The combination of Metadynamics and extended Lagrangian Adaptive Biasing Force (Meta-eABF) is an advanced computational method used to efficiently explore conformational landscapes and calculate free energy profiles along predefined CVs^21,39^. Since our goal was to sample the slow transitions between symmetric and asymmetric sTNF conformations, we analyzed symmetric (1TNF) and asymmetric (7JRA) crystal structures to identify features that would effectively distinguish these two states and thus would serve as CVs to drive Meta-eABF simulations. Our analysis identified three candidate distances as the CVs to use for accelerating the structural transition between monomers A and C. S1, S2, and S3 are defined as the distances between the center of masses of Cα atoms (CoM) of the following residues (see **Fig. 2c**):

S1: distance between CoM of residues 130 131 132 200 201 202 of subunits A and C.
S2: distance between CoM of residues 169 170 171 172 173 174 193 194 195 196 197 198 of subunit A and residues 136 137 138 139 140 192 193 194 195 226 227 228 of subunit C
S3: distance between CoM of residues 151 152 153 173 174 175 of subunit A and residues 140 141 142 190 191 192 of subunit C.

According to these definitions, S1 measures the CoM distance between the same segments of subunits A and C, whereas S2 and S3 are measuring CoM distances between different segments of subunits A and C. As seen from the table in **Fig. 2d**, all three distances are significantly larger in the asymmetric state compared to the symmetric conformation. We note that another member of the TNF family, namely CD40L^13^, has been crystallized in symmetric and asymmetric conformations (PDB IDs 1ALY and 3LKJ). The three proposed CVs are transferable to CD40L and likely to other TNF proteins that share the same trimeric assembly as TNF and CD40L^13^. Thus, we used S1, S2, and S3 CVs for the Meta-eABF simulations.

For the meta-eABF simulations, we used OpenMM 7.7 patched with PLUMED 2.8.1^40^. The system was heated from 0 to 310 K in 0.12 ns time intervals, followed by a production phase at 310K. We additionally used hydrogen mass repartitioning^34^ which allowed integration timestep of 4 fs, and Langevin thermostat with friction of 1.0 ps^-1^. For the sTNF and mTNF constructs, we performed 2 independent meta-eABF simulations each, yielding cumulative sampling times of 4 μs and 2 μs, respectively. The eABF bias was applied to S1, S2 and S3 distances. To limit the exploration of the phase space, we chose the following ranges for S1, S2 and S3: [17,26] Å, [7,15] Å, and [9, 15] Å. Well-tempered Metadynamics was performed on the fictitious variables S1_fict, S2_fict and S3_fict. Details about real and fictitious variables of the Meta-eABF method can be found in ref^21^.

To generate the free energy surface (FES) in the space of S1, S2 and S3 CVs, we integrated the force bias obtained from each meta-eABF simulation using the Poisson integrator in ref^41^. The calculations were performed on the rectangular grid of 91×81×61 nodes spanning S1 ∈ [17.0, 26.0] Å, S2 ∈ [7.0, 15.0] Å, S3 ∈ [9.0, 15.0] Å, with 0.1 Å spacing. To ensure that the reported free energies rest on adequately sampled regions of the CV space rather than a small number of sparsely visited outlier nodes, the threshold of ≥ 100 on node counts was used to mask all free-energy calculations. As shown in **Figs. S2-S3**, this cut-off retained >93% of the sampled domains for all the simulated systems.

##### Time-lagged independent component analysis (tICA) of the unbiased MD simulations

As in previous studies^42–45^, we used tICA^46^ to perform dimensionality reduction of the unbiased MD simulations. Briefly, two different covariance matrices are constructed from the MD simulation trajectories: a time-lagged covariance matrix (TLCM) **C**^TL^(τ) = <**X**(t)**X**^T^(t+τ)>, and the standard covariance matrix **C** = <**X**(t)**X**^T^(t)>, where **X**(t) is a vector (function of the system’s positions) at time t, τ is the lag-time of the TLCM, and the symbol <…> denotes the time average. To identify the slowest reaction coordinates of the system, the following generalized eigenvalue problem is solved: **C**^TL^ **V** = **C V** Λ, where Λ and V are the eigenvalue and eigenvector matrices, respectively. The eigenvectors corresponding to the largest eigenvalues define the slowest reaction coordinates. These reaction coordinates depend on the choice of data vector X, i.e., the choice of the input features. Here, to define the tICA space, we considered two sets of input features. The first set quantified membrane binding of the mTNF ECD and contained z-directional (vertical) distances between the phosphorus atoms of the extracellular membrane leaflet and the Cα atoms of the following set of residues: R107/R108, R120, and R207. The second set of input features quantified intra-ECD conformational fluctuations and contained the S1, S2, and S3 distances described above. To facilitate structural analysis of the tICA data, the 2D tICA space was discretized into 50 microstates (referred throughout simply as “states”) using automated clustering k-means algorithm.

#### Structural analysis

To quantify the structural similarity between a simulation trajectory and the crystal structure of the symmetric (PDB id 1TNF) or asymmetric state (PDB id 7JRA), we calculated the Root Mean Square Deviation (RMSD) in the following way: (i) we aligned the trajectory and crystal structure on the Cα atoms of the β sheets in subunit C; (ii) we calculated the RMSD between subunit A of the trajectory and the symmetric structure.

## RESULTS

### Membrane binding of mTNF ECD is facilitated by positively charged residues on ECD

To study membrane binding of mTNF ECD, we carried out extensive unbiased atomistic MD simulations. As detailed in the Methods section, the simulations were started from 20 mTNF models differing in their ECD position with respect to the membrane surface, the conformation of the ECD, i.e., whether it was initially in the symmetric or asymmetric conformation (**Fig. 2c-d**) and the conformation of the linker region (see Methods and **Fig. S1b**). Each of these 20 systems was simulated with unbiased MD for 1 μs.

During the dynamics, we observed multiple events of ECD binding to the lipid membrane. As shown in **Table 1** and **Fig. 3**, we found two different membrane binding modes. In the first, Mode 1, the ECD engaged with the membrane primarily through several positively charged residues on one of its subunits (**Fig. 3a-c**). These residues include R78/R82, R107/R108, R120, and R207 (see also **Figs. S4-S7**). This binding mode, observed for all three subunits of mTNF in different trajectories (**Table 1** and **Fig. 3a-c**), is characterized by an overall tilting of the ECD domain with respect to the membrane normal axis. As shown in the time-evolution traces of the ECD tilt angle (**Fig. S8**, the trajectories marked by “x”), Mode 1 is characterized by an average tilt angle of 54±13 degrees. Similar conformations of the ECD, with an average tilt angle of ∼60 degrees, have been reported in a previous simulation work^22^. In the second binding mode, Mode 2, the ECD associated primarily through residues R78 and R82 from all three subunits (**Fig. 3d** and **Figs. S4-S7**). In this binding mode, the ECD domain remains relatively vertical, with an average tilt angle of 18±7 (**Fig. S8**). In both, Mode 1 and Mode 2, the protein engaged mainly with POPC and SM lipids (**Fig. S9**) which are a majority lipid species on the EC membrane leaflet.

**Figure 3:**
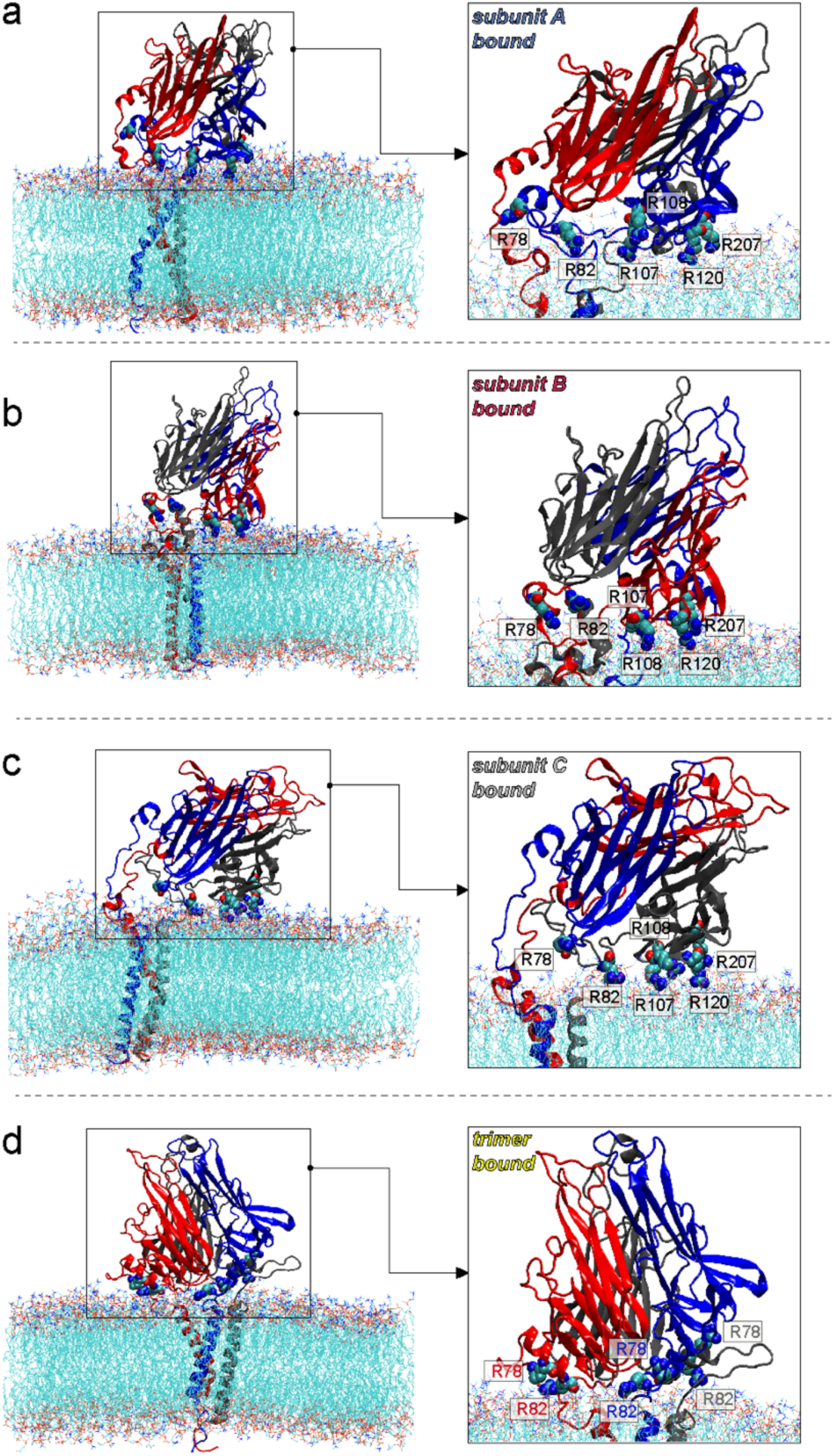
Membrane binding modes of mTNF ECD. **(a-c)** Membrane binding Mode 1 established by the ECD of subunit A (a), B (b), and C (c) in MD simulations. **(d)** Membrane binding Mode 2 established by the entire ECD trimer in MD simulations. In each panel, mTNF-membrane complex is shown on the left and the detailed views of the respective binding poses are shown in the right rectangles. In all the images, only the non-hydrogen atoms of membrane lipids are depicted and the solution atoms (waters, ions) are removed for clarity. The protein is colored as in Fig. 2, with selected residues participating in membrane binding highlighted in space-fill and labeled.

**Table 1:** List of unbiased MD simulations of mTNF.

| **System** | **Replicate** | **Membrane Binding Mode** |
|---|---|---|
| mTNF1<sup>Sym</sup> | 1 | no binding |
|  | 2 | monomer A - R207, R120, R107/R108, R82/78 |
|  | 3 | monomer B - R207, R120, R107/R108, R82/78 |
|  | 4 | no binding |
|  | 5 | monomers A, B, C - only R82/78 |
| mTNF2<sup>Sym</sup> | 1 | no binding |
|  | 2 | monomer B - R207, R120, R107/R108; monomer A - R82/78 |
|  | 3 | no binding |
|  | 4 | no binding |
|  | 5 | no binding |
| mTNF1<sup>Asym</sup> | 1 | no binding |
|  | 2 | monomer C - R207, R120, R82/78 |
|  | 3 | no binding |
|  | 4 | no binding |
|  | 5 | monomers A, B, C - R82/78 |
| mTNF2<sup>Asym</sup> | 1 | no binding |
|  | 2 | monomer C - R207, R120, R107/R108, R82/78 |
|  | 3 | monomer A - R207, R120, R107/R108, R82/78 |
|  | 4 | no binding |
|  | 5 | no binding |

As shown in **Table 1**, the system dynamically converged to Mode 1 binding irrespective of the starting positioning of the ECD domain (i.e., the initial linker conformation, mTNF1 or mTNF2). Importantly, Mode 1 was more prominent than Mode 2, as we found Mode 1 fully formed in 4/20 replicates and partially formed in additional 2 replicates. Specifically, in replicate 2 of mTNF2^Sym^, R78/R82 from another subunit is engaged with the membrane, while the binding mode in replicate 2 of mTNF1^Asym^ lacks R107/R108 engagement with the membrane. Mode 2 was detected in only 2/20 replicates.

To assess the dynamic stability of the two binding modes, we computed the fraction of simulation time that any Arg residue among those listed above were in contact with the membrane. To this end, we defined a contact cutoff of 10 Å and calculated the percentage of frames in which the closest Arg Cα–P distance was below the cutoff (**Figs. S4–S7**). This analysis revealed that, once the ECD established interactions with the membrane, it remained in contact with the lipids for 67-98% of the trajectory frames (depending on the simulation replicate), suggesting that the two binding modes were stable on the simulation timescales.

### ECD-membrane interactions and protein conformational changes

To better describe the ECD-membrane interactions and concomitant conformational changes of the ECD, we carried out tICA dimensionality reduction of the MD data. As described in the Methods section, we used the following tICA input features: three distances quantifying position of residues R107/R108, R120, and R207 in mTNF ECD with respect to the membrane surface, and distances S1, S2, and S3 quantifying intra-ECD rearrangements during symmetric-to-asymmetric transitions (see **Fig. 2c-d**).

**Fig. 4a** shows the projections of the MD trajectories onto the 2D tICA space defined by the first two tIC vectors. **Fig. S10a** indicates the locations of the initial four structures (mTNF1^Sym^, mTNF2^Sym^, mTNF1^Asym^, and mTNF2^Asym^) on the 2D tICA space. The first two tIC vectors capture 98% of the fluctuations in the system (**Fig. S10c**) and they describe the two slowest molecular processes. Specifically, tIC1 mostly includes contributions from the distances between R107/R108, R120, and R207 and the membrane surface (**Fig. S10d**), indicating that tIC1 encodes information about the ECD membrane binding. Indeed, the MD trajectories in which the ECD binds to the membrane progress in time through the tICA space from left to right direction (**Fig. S11**, see the replicates marked by “x” symbol). On the other hand, tIC2 primarily encodes S1, S2, and S3 distances, suggesting that tIC2 describes the internal ECD slow dynamics.

**Figure 4:**
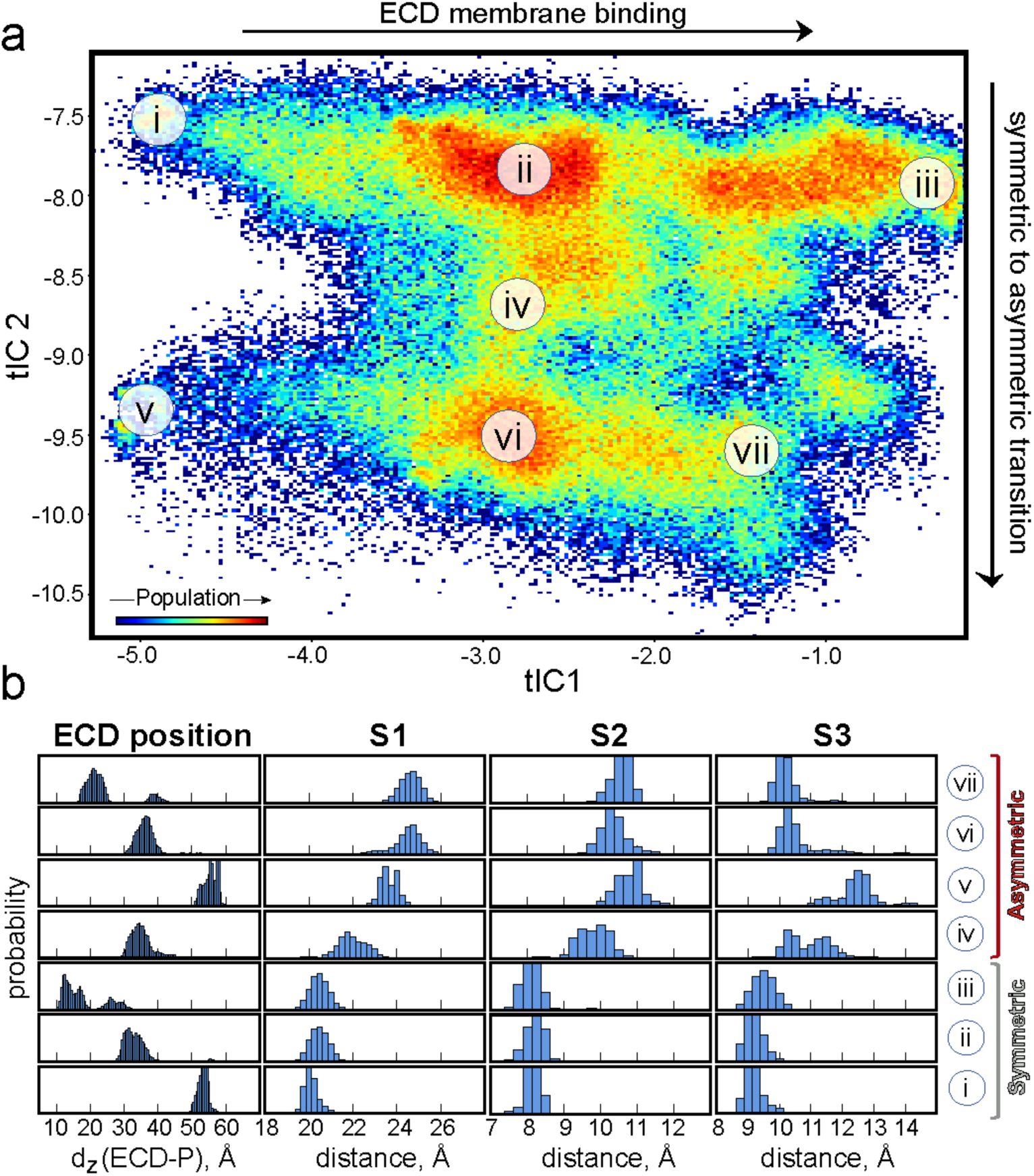
Conformational sampling of mTNF ECD in MD simulations. **(a)** Projection of all the trajectory frames from the unbiased MD simulations onto the 2D tICA space spanned by the first two tIC vectors. The color map identifies the populations distribution of the different states of the system in this tICA space, and the locations of selected states on the 2D space are indicated by Roman numerals. tIC1 encodes dynamics of the ECD membrane binding, whereas tIC2 describes symmetric-to-asymmetric transitions within the ECD. **(b)** The histograms quantifying the position of the ECD with respect to the membrane and the ECD structural features for each of the selected states from panel *a*. From left to right the histograms show: the vertical (z-directional) distance between the ECD center-of-mass and the center-of-mass of the lipid phosphorus atoms in the extracellular membrane leaflet; S1, S2, and S3 distances that quantify various degrees of asymmetry within the ECD. States i-iii are characterized by the symmetric ECD structure, whereas states iv-vii consist of asymmetric ECD conformations (see also Fig. 2c).

To gain a deeper understanding of the molecular mechanism, we discretized the tICA space into 50 states (see Methods and **Fig. S10b**) and calculated several structural properties in each state (**Fig. 4** and **Fig. S12**). When traversing the tICA space from left to right, the ECD gradually approaches and ultimately binds the lipid membrane. This is reflected in decreased vertical distance between the ECD center-of-mass and the membrane phosphate plane (see changes in “ECD position” between states i-iii and v-vii in **Fig. 4b**). Consistent with this trend, the states in the rightmost part of the tICA space are characterized by relatively short distances between the selected basic residues (R82, R107/R108, R120, and R207) and the membrane phosphate plane (see changes in **Fig. S12a-c** histograms between states 1-5 and 10-14).

### The mTNF ECD domain transitions between symmetric and asymmetric conformations

We next examined structural changes within ECD during the unbiased simulations. To this end, we compared histograms of the S1, S2, and S3 distances in different states on the tICA space (**Fig. 4b** and **Fig. S12d**). The analysis revealed that in the states located in the upper part of the space (tIC2 > -8.0, i.e., states i-iii in **Fig. 4b** or 1-5 in **Fig. S12d, S12g**) S1, S2, and S3 fluctuate around the values measured in the symmetric state (i.e., S1∼20 Å, S2∼8 Å, and S3∼9 Å, see **Fig. 2d**). However, for the states in the lower part of the tICA space (tIC2 < -8.0), S1, S2 and S3 sample values that are no longer consistent with the Symmetric^X-ray^ structure (**Fig. 2b-c**). Rather, we observe the ECD dynamically sampling different asymmetric states. Specifically, the conformations that are most similar to the asymmetric crystal structure belongs to state v in **Fig. 4** (or state 10 in **Fig. S12d, S12g**). The conformations in this state are characterized by relatively larger values of S1, S2, and S3 variables (see histograms in **Fig. 4b**), consistent with those from the Asymmetric^X-ray^ structure (S1∼24Å, S2∼11Å, and S3∼12Å, see **Fig. 2d**). We also observed asymmetric conformations in which S1, S2, and S3 variables assume intermediate values between the ones measured in Symmetric^X-ray^ and Asymmetric^X-ray^ structures. For example, the ensemble of conformations belonging to state iv in **Fig. 4** are characterized by S1∼22 Å, S2∼10 Å, and S3∼11 Å. States vi-vii in **Fig. 4** contain conformations with somewhat larger S1 (∼25 Å) but shorter S2 (∼10.5 Å) and S3 (∼10.5Å), compared to the crystallographic asymmetric structure.

Taken together, the different conformations of the ECD observed in our simulations can be separated into four distinct structural classes as shown in **Fig. 5**. The symmetric state akin to the Symmetric^X-ray^ structure originates from regions i-iii of the tICA landscape depicted in **Fig. 4**. Asymmetric states 1, 2, and 3 depicted in **Fig. 5** originate from regions iv, v, and vi-vii in **Fig. 4**, respectively, with Asymmetric 2 being most similar to the Asymmetric^X-ray^ structure. The average RMSD of the asymmetric classes 1, 2, and 3 with respect to Symmetric^X-ray^ (see Methods) is 4.8 Å, 5.5 Å and 6.4 Å, respectively.

**Figure 5:**
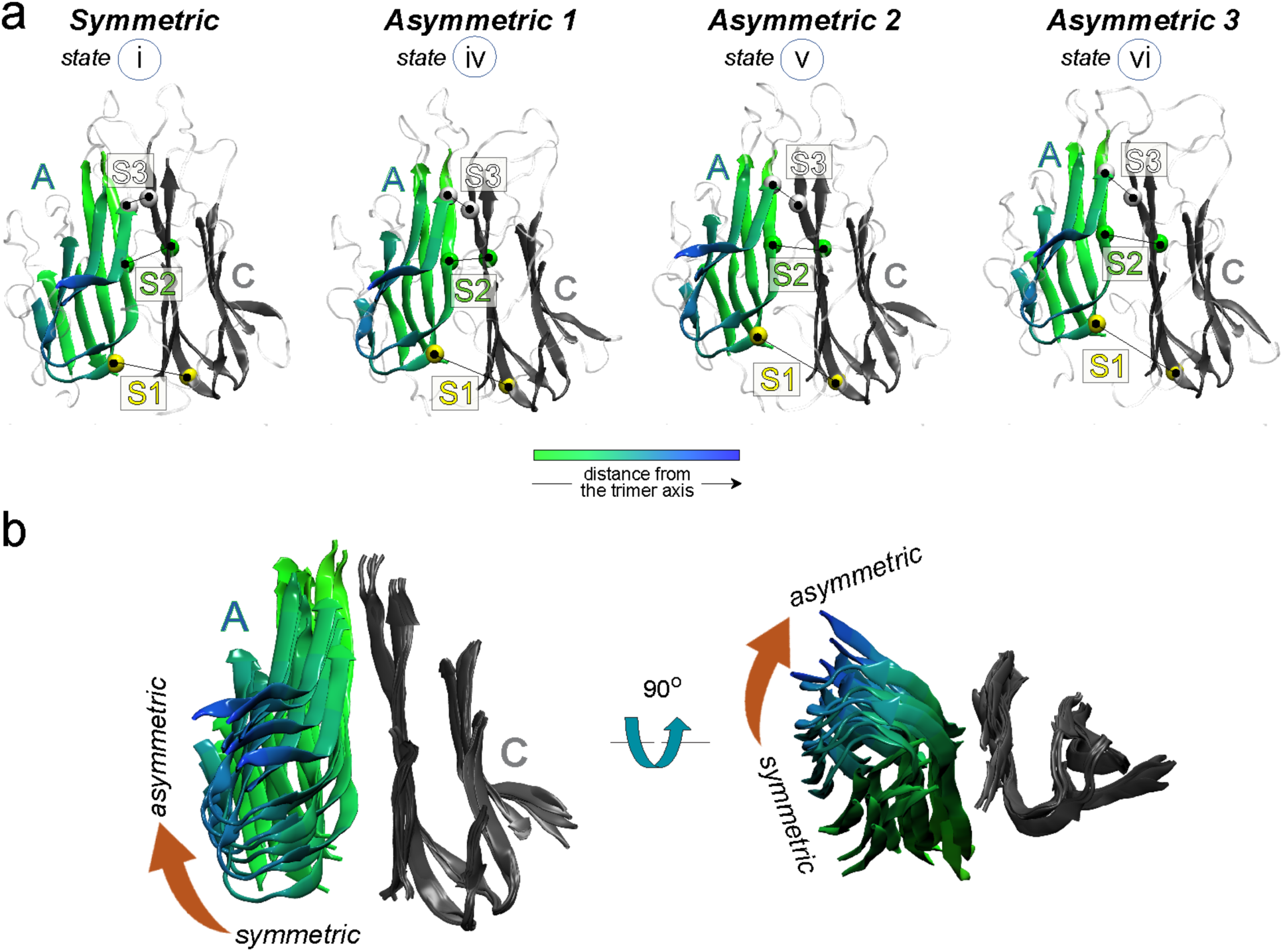
Structural characteristics of the asymmetric states of mTNF ECD from MD simulations. **(a)** Representative structures of the ECD from states i, iv, v, and vi of the tICA space shown in Fig. 4. These are referred throughout as Symmetric, Asymmetric 1, Asymmetric 2, and Asymmetric 3 states, respectively. Asymmetric 2 is the most similar to the asymmetric crystal structure 7JRA. Only subunits A and C are shown for clarity (subunit B is omitted). Subunit C is colored grey and the atoms on subunit A are colored according to their distances from the trimer vertical axis with blue shades corresponding to the most distant regions. **(b)** Superposition of all 4 structures from panel *a* illustrating gradual outward and upward movement of subunit A with respect to subunit C.

To gain additional molecular insights regarding intra-ECD rearrangements during the conformational transitions, we calculated the distance *d*_198−224_ between the Cα atoms of residues G224 and G198 on two adjacent TNF monomers. We chose this specific distance because, akin to the movement of an opening door, *d*_198−224_ increases from 3.8 Å (PDB ID 7JRA) to 9 Å (PDB ID 1TNF). Indeed, as illustrated in **Fig. S12e**, the *d*_198−224_ values are distributed between 4 and 6Å for the asymmetric conformations and between 8 and 12 Å, for the symmetric conformations. Our analysis reveals that this increase in *d*_198−224_ during the asymmetric-to-symmetric transition is accompanied by the loss of several interactions between the GH loop of subunit C (residues 221 to 225) and the E strand of subunit A, and formation of the new ones between the same loop and the F strand of subunit A (**Fig. S12f**).

### The mTNF ECD binds to the membrane predominantly in the symmetric conformation

We next sought to establish whether the ECD binds the membrane in a specific conformation. To this end, we tracked the dynamic progression of the MD trajectories on the tICA space (**Fig. S11**. We found that from the 8 simulations in which the ECD associated with the membrane (indicated by the “x” symbol in **Fig. S11**), in 5 replicates the membrane-bound ECD was in the symmetric state, i.e., these trajectories projected to the upper part of the tICA space. Interestingly, from the four runs in which Mode 1 binding was observed (replicates 2-3 in the mTNF1^Sym^ set and replicates 2-3 in the mTNF2^Asym^ set), three showed membrane binding of the symmetric ECD (the exception was replicate 3 from the mTNF2^Asym^ set). The ECD in replicate 2 from the mTNF2^Asym^ set underwent full asymmetric-to-symmetric transition upon membrane binding. To better understand the molecular basis for this, we performed detailed analysis of this replicate by quantifying interactions involving the basic residues on the ECD implicated in the membrane binding (R82, R107, R108, R120, and R207).

### Asymmetric-to-symmetric transition leads to ECD conformational changes that facilitate membrane binding

Our analysis of simulation replicate 2 from the mTNF2^Asym^ set uncovered how the asymmetric-to-symmetric transition between subunits A and C can favor the binding between subunit C and the membrane. **Fig. 6** shows structural representations and dynamic changes along the interface between subunits A and C of the ECD by plotting the time evolution of the *d*_198−224_ distance (see also **Fig. S12f**). At the initial stages of the simulation, *d*_198−224_ samples relatively short distances. But as the asymmetric-to-symmetric transition proceeds, *d*_198−224_ gradually increases towards symmetric-like values.

**Figure 6:**
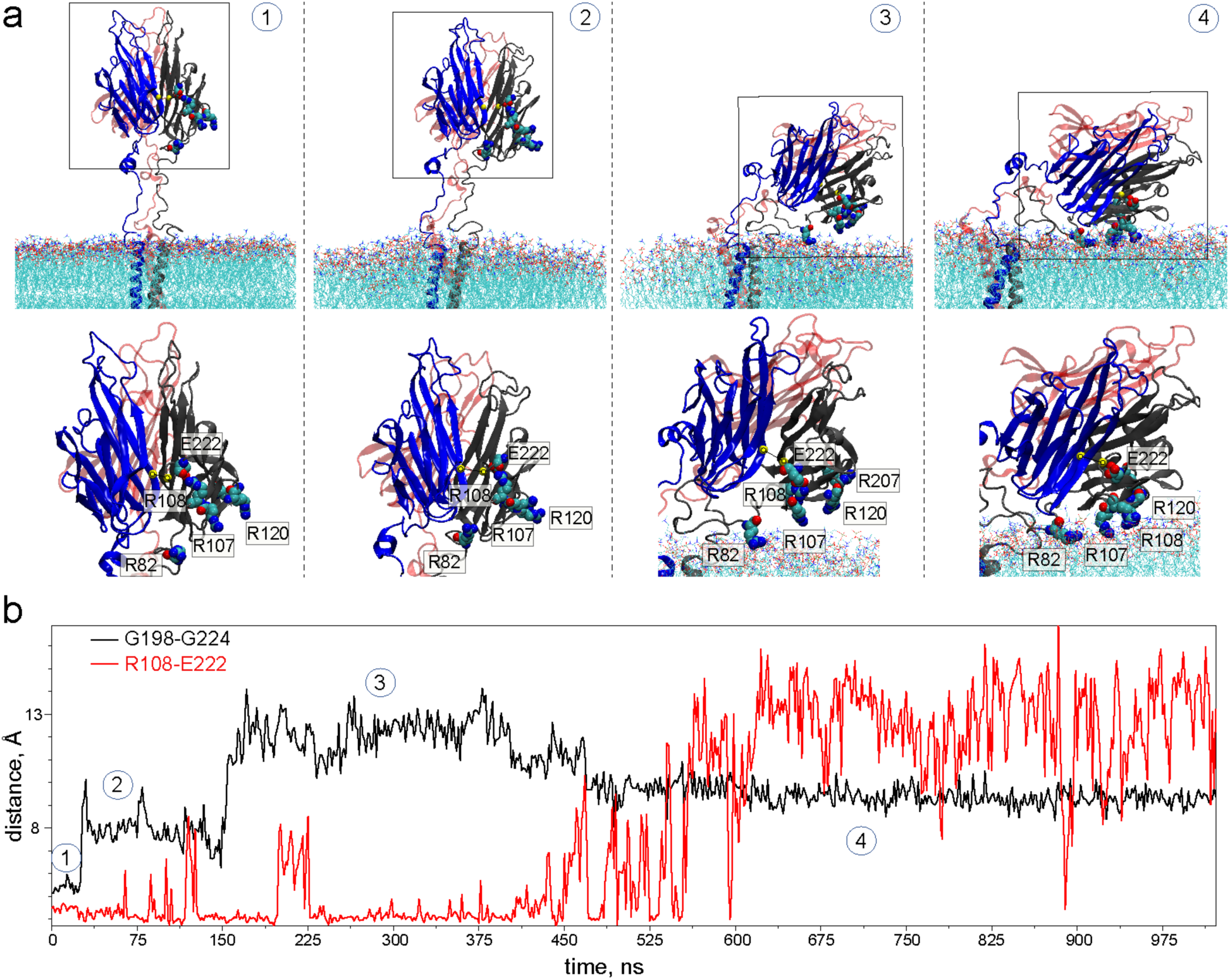
Symmetrizing of mTNF ECD upon membrane binding. **(a)** Structural representation of the key events during subunit C membrane binding observed in MD simulations of replicate 2 from the mTNF2^Asymm^ set. Highlighted are changes in R108-E222 interactions and local structural rearrangement as subunit C establishes Mode 1 membrane binding. The bottom panels show zoomed-in views of the parts of the system marked by rectangles. Relevant residues are shown in space-fill and are labeled. **(b)** Time-evolution of G198-G224 (black) and R108-E222 (red) distances in the simulation from panel *a*. The trajectory points representing the 4 snapshots in panel *a* are indicated.

This movement has important implications for how subunit C can establish Mode 1 membrane binding. Specifically, in the starting structure (Asymmetric^X-ray^), E222 and R108 are buried within the subunit C and form a salt bridge (**Fig. 6**, conformation 1). As the 221-225 loop moves away from the F beta strand of subunit A, during the asymmetric-to-symmetric transition, the increased flexibility of E222 leads to breaking of the salt bridge between E222 and R108 (**Fig. 6**, conformations 2-3). Once the E222-R108 bond is broken, the R108 sidechain becomes solvent exposed, which allows it to insert into the membrane and stabilize Mode 1 binding (**Fig. 6**, conformation 4).

The analysis of E222-R108 interactions across all our simulations (**Fig. S13**) revealed that this interaction continues to be predominant in subunit C but is rarely seen in subunits A and B. Importantly, in the replicate in which subunit C binds membrane in Mode 1, but without the involvement of R107/R108 (replicate 2 from mTNF1^Asym^ set), E222-R108 interactions are relatively preserved (in ∼80% of the trajectory frames). In all other simulations in which full Mode 1 binding is established (i.e., with R107/R108 also engaged with the membrane), E222-R108 interactions are either not present or transient (**Table 1** and **Fig. S13**). Taken together, these results suggest that breaking of the E222-R108 salt bridge, during the asymmetric-to-symmetric transition, leads to conformational changes in the ECD that promote Mode 1 membrane binding.

### The membrane energetically stabilizes symmetric mTNF states

The data from the unbiased MD simulations presented above suggest that the symmetric conformation of the ECD is predominant in mTNF and is further favored when the ECD binds to the membrane through Mode 1. To test this hypothesis quantitatively, we used biased meta-eABF MD protocol using S1, S2 and S3 as CVs (see Methods). With these simulations, our goal was to assess energetic feasibility of the symmetric vs asymmetric conformations when the mTNF is engaged in Mode 1 membrane binding. Specifically, we sought to calculate the free energy differences between the symmetric and asymmetric states in the mTNF interacting with the membrane via Mode 1 and compare these differences to those in sTNF. To this end, we considered two protein models: the full-length membrane bound mTNF taken from the unbiased MD simulations of replicate 2 of the mTNF2^Asym^ set (**Fig. 6**, ECD engaged with the membrane in Mode 1), and sTNF. For each of these constructs, 2 independent microsecond-scale meta-eABF simulations were performed (see Methods). During these simulations, we observed broad coverage of the 3D conformational space of the CVs (**Figs. S14-S15**).

We first analyzed ECD-membrane interactions in the mTNF meta-eABF simulations to confirm that the ECD was stably bound to the lipids. Indeed, as shown in **Fig. S16**, the Mode 1 binding was maintained throughout these meta-eABF simulations.

To quantify energetics from the meta-eABF trajectories, we next calculated the FES in the 3D space of the three CVs for mTNF (2 replicates) and sTNF (2 replicates). The four FESs, depicted in **Fig. 7a-b**, show low energy minima characteristic of the symmetric ECD conformation (see table in **Fig. 7** and the histograms in **Fig. 4**). The two sTNF FESs also show a diffuse energy landscape of low energy asymmetric conformations. In contrast, for the mTNF construct the landscapes largely consist of a single, well-defined energy minimum corresponding to the symmetric state ensemble, which appears to be separated from the other regions of the FES by a larger energy barrier compared to sTNF. The extensive sampling of the S1/S2/S3 (histograms in **Fig. S2-S3, S14-S15**) confirms that both symmetric and asymmetric free energy minima were frequently visited during the meta-eABF simulations.

**Figure 7:**
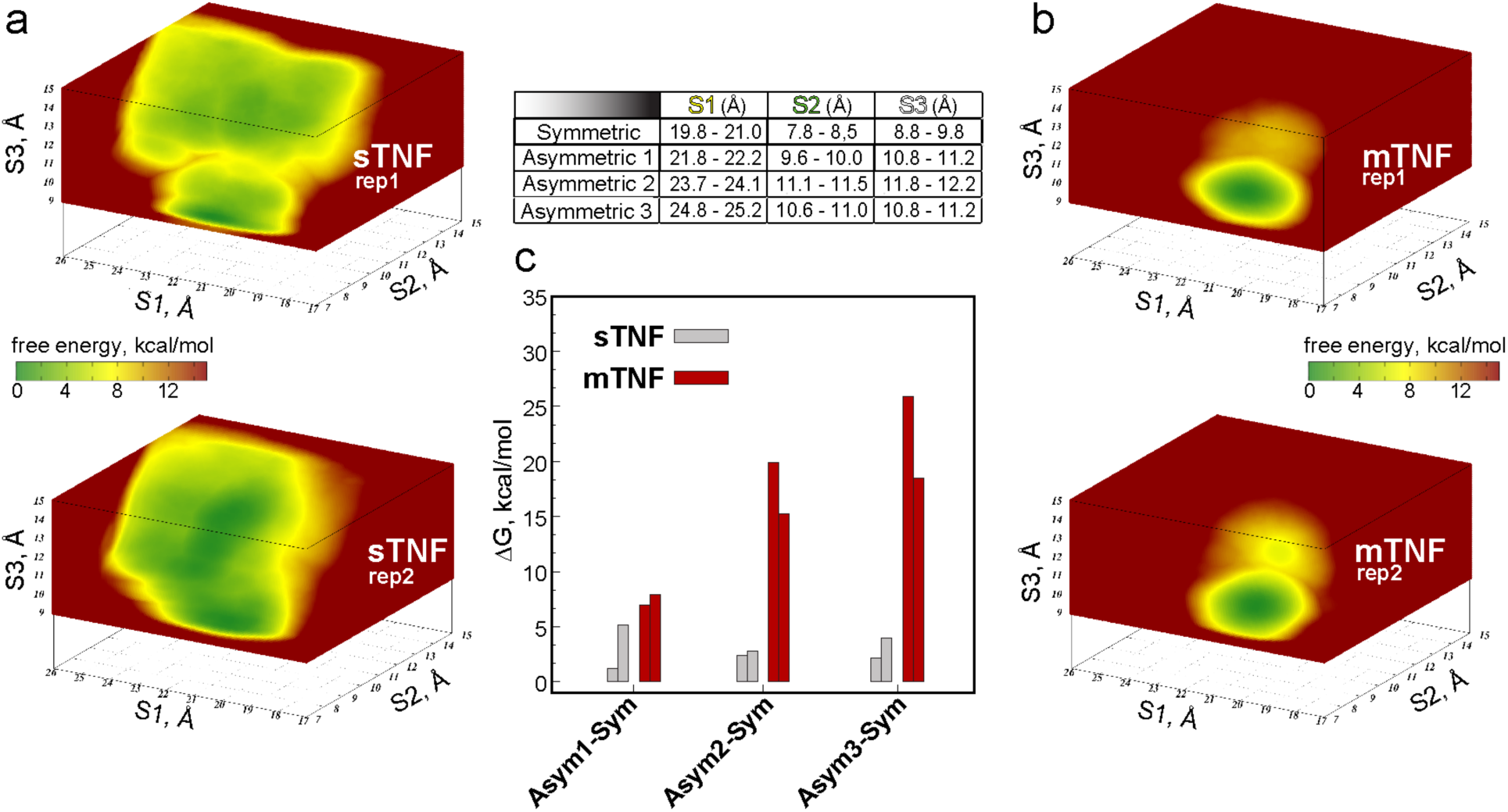
Membrane binding stabilizes symmetric conformations in mTNF. (**a**-**b**) FES in the 3D space of S1, S2, and S3 variables for two independent replicates of sTNF (a) and mTNF (b) systems from meta-eABF MD simulations. The landscape is colored according to the color bar. (**c**) The free energy difference between the main Asymmetric states 1, 2 and 3 and the Symmetric state (Δ*G_Asym_*_1−*Sym*_, Δ*G_Asym_*_2−*Sym*_, and Δ*G_Asym_*_3−*Sym*_) were obtained from the FESs, and reported for sTNF (grey bars) and mTNF (red bars) systems. Each bar correspond to a simulation replicate. The locations of the symmetric and 3 asymmetric states on the FES were defined according to the S1, S2, and S3 value ranges given in the table.

To calculate the free energy of each TNF state from the 3D FESs, we defined S1, S2, S3 ranges corresponding to the 3 asymmetric states found in the unbiased MD simulations (see table in **Fig. 7** and histograms in **Fig. 4**). We then calculated the free energy of these populations from the FESs of sTNF and mTNF (see **Fig. S17** for convergence analysis). **Fig. 7c** shows that the sTNF asymmetric states 1, 2, and 3 are characterized by ∼2-5 kcal/mol higher free energy compared to the symmetric state. This energy difference is in good agreement with previous estimates^9^. The energy difference between the asymmetric and symmetric states in the mTNF system is significantly higher compared to that in sTNF. Thus, as shown in **Fig. 7c**, in mTNF, the Asymmetric states 1, 2, and 3 are characterized by 4.3 ± 2.0 kcal/mol,15.0 ± 2.3 kcal/mol, and19.1 ± 3.8 kcal/mol higher free energies, respectively, than the symmetric state. Because the FES we obtained is based on the CVs that describe symmetric-asymmetric transitions driven by the pair of the TNF trimer from which one of the subunits is membrane bound, the predicted energy values likely represent the upper bound of the free energy for these transitions. Overall, our results suggest that the membrane environment stabilizes the mTNF symmetric state compared to asymmetric conformations. This conformational selectivity on the membrane surface is likely driven by the combination of entropic effects arising from spatial confinement^47^ due to steric effects stemming from the bilayer and the favorable interactions between TNF basic residues and membrane lipids^48,49^.

## DISCUSSION

TNF plays a central role in inflammatory pathways and remains a major therapeutic target for chronic diseases such as rheumatoid arthritis, inflammatory bowel disease and psoriasis^1,50^. Despite decades of therapeutic success targeting TNF, a detailed mechanistic understanding of its structural and energetic behavior has remained incomplete, limiting efforts to differentially modulate soluble and membrane-bound forms. Historically, TNF inhibition has been achieved primarily through biologics that act by competing with receptor binding. More recently, the conformational plasticity of TNF has been leveraged to design small molecules that bind and inhibit TNF activity through an allosteric mechanism^12^. However, this approach remains challenging, and only two such small molecules have reached clinical trials to date^16,51^.

The soluble form of TNF, sTNF, exists in a dynamic equilibrium between symmetric and asymmetric conformations^8,9^. Experimental studies have provided complementary structural descriptions of this conformational heterogeneity: DEER measurements have been used to derive models of asymmetric states sampled in the apo sTNF^8^, whereas X-ray structures have captured asymmetric conformations stabilized by small molecules^9^. In the latter case, binding of small molecules to the inter-subunit cavity located at the trimer center disrupts the trimer symmetry and consequently prevents full receptor binding^9^. Acidic conditions can also induce an asymmetric, receptor-incompetent form^52^, further suggesting that sTNF conformational landscape is heterogeneous.

The transmembrane form of TNF, mTNF, has recently become an attractive target because it is involved in inflammation and immune response by acting as a cell-to-cell signaling molecule^1^. However, in contrast to sTNF, the structural and dynamic properties of mTNF have remained elusive, largely due to the experimental challenges of characterizing membrane proteins.

In this work, we employed a combination of molecular modeling of the full-length mTNF, atomistic MD simulations and advanced analysis techniques to characterize the conformational landscape of mTNF. Our analysis of multi-microsecond unbiased MD trajectories identified two dominant processes: (1) the asymmetric-to-symmetric structural transition of the ECD and (2) the association of the ECD with the membrane surface. In the unbound state, the ECD populates multiple asymmetric intermediates along the transition pathway toward the symmetric trimer. Upon membrane association, however, conformational heterogeneity is markedly reduced, and the symmetric state becomes predominant, indicating that membrane engagement spatially restricts ECD motions and thermodynamically stabilizes the symmetric conformation relative to the asymmetric one.

Detailed inspection of the unbiased MD simulation trajectories highlighted two preferred membrane-binding modes. In the more frequently observed Mode 1, the ECD is in the symmetric state and adopts a ∼54 degrees tilt relative to the membrane normal, stabilized mostly by electrostatic contacts involving residues R78–R82, R107, R108, R120, and R207 with the lipid headgroups. The conformational transition of the initially solvent-exposed asymmetric ECD, which leads to Mode 1, begins with the ECD positioned far from the lipid membrane and with a fully formed salt bridge between residues E222 and R108 (see **Fig**. **6** and **Fig. 8**). The key dynamic event preceding membrane engagement is the disruption of the E222–R108 salt bridge, which facilitates the concurrent structural transition of the ECD from asymmetric to symmetric. After the salt bridge breaks, residue R108 establishes stable interactions with the membrane lipid headgroups, remaining in contact for 83% of the simulation time (average over six independent trajectories). In Mode 2, the mTNF trimer adopts an orientation nearly perpendicular to the membrane surface. In this conformation, residues R78 and R82 from each monomer form long-lived electrostatic interactions with the lipid headgroups, anchoring the trimer to the membrane. The effect that the membrane elicits on mTNF is consistent with prior studies on different membrane-anchored proteins, whose ectodomains either reorient or undergo structural transitions upon membrane engagement ^53–56^.

**Figure 8:**
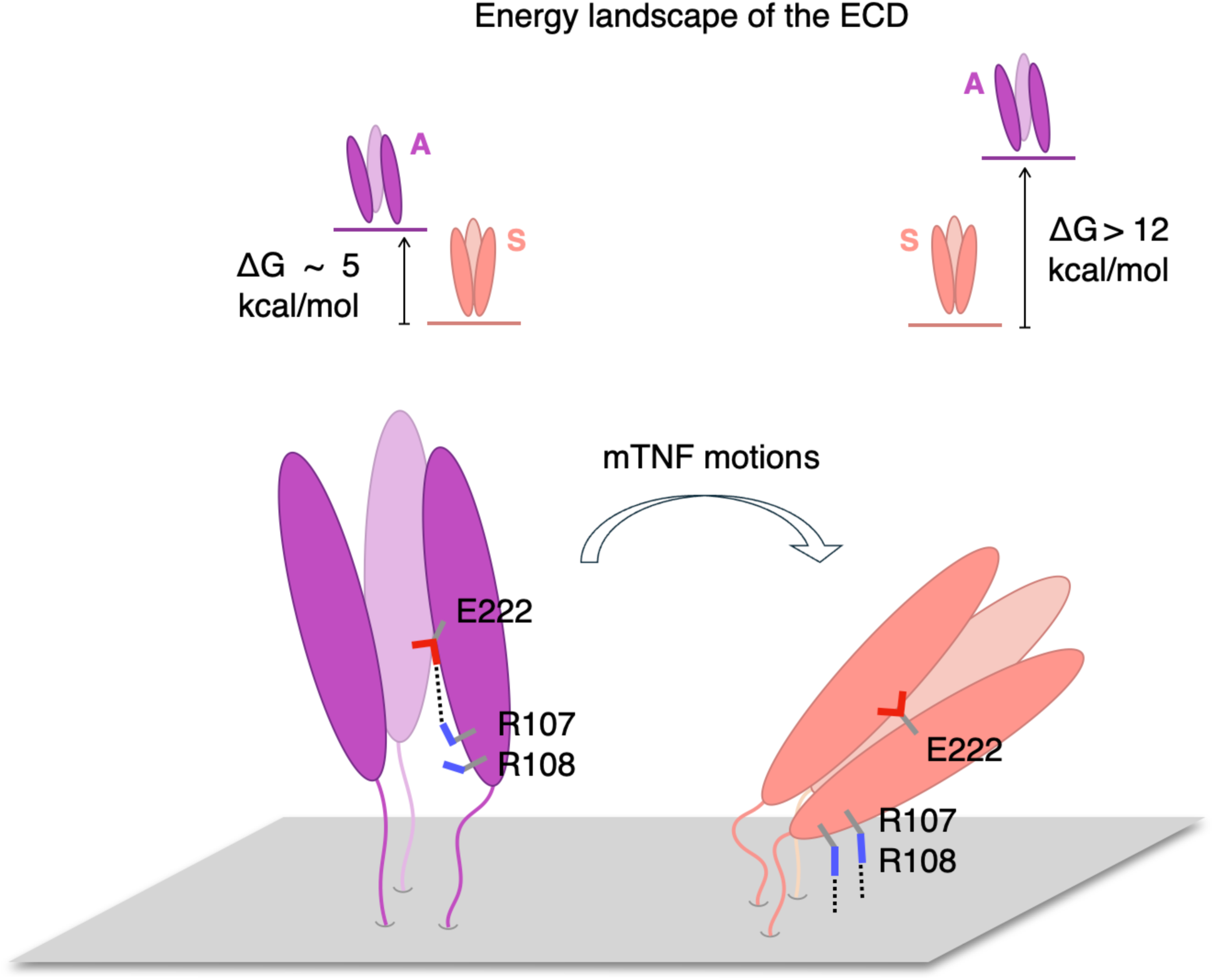
Conformational and energetic consequences of mTNF - membrane association from computer simulations. Illustration of the mTNF concerted motions explored by our simulations: salt bridge E222-R107 disruption, intra-ECD asymmetric (A) to symmetric (S) conformational change and ECD-membrane association. S → A free energy differences for the two mTNF states are shown in the upper panels, supporting the conclusion that the membrane stabilizes the symmetric conformation.

ECD-membrane interactions have been reported for other TNF superfamily members such as TRAIL and GITRL^14,23^ or the complement component C1q^57^. Moreover, studies using engineered mTNF constructs showed that mTNF can assemble into clusters with defined three-dimensional orientations, which strongly enhance TNFR2 assembly and activation^58^. The above observations suggest that ECD–membrane contacts may represent a conserved mechanism to regulate downstream processes such as the association of TNF with its cognate receptor ^4,19^.

The limited timescales accessible to unbiased MD simulations precluded an accurate estimate of the TNF free energy landscape. To overcome this limitation and accelerate the symmetric ↔ asymmetric transitions in both sTNF and mTNF models, we employed meta-eABF simulations^39^. Our goal was to estimate the energetic stability of the symmetric state relative to the asymmetric states when mTNF engages with the membrane in Mode 1 binding and compare it to the energy differences between these states in sTNF. In sTNF, we estimated free energy differences between asymmetric and symmetric states on the order of ∼2-5 kcal/mol, consistent with previous experimental and simulation studies^8,9^. This difference is larger in mTNF suggesting that binding of the ECD to the lipid membrane stabilizes mTNF symmetric state compared to sTNF. The stabilization of the symmetric state by the membrane, together with the stable ECD–membrane association, may provide a mechanistic basis for the 10-to 100-fold reduction in activity of certain biologics toward mTNF^17^. This reduction is likely multifactorial, with membrane effects acting as one contributing factor, as suggested by the present work. Indeed, by engaging with the membrane in the symmetric conformation as described here, the ECD domain of mTNF may become inaccessible to biologics due to a steric obstruction caused by the juxtaposition of the lipid membrane. To evaluate this possibility, we considered, as a typical example of biologics-bound TNF, the X-ray structure of the ECD trimer bound to 3 golimumab molecules (PDBID: 5YOY^59^). We then performed structural alignment of the ECD from this system onto the ECD of the membrane-bound mTNF and calculated the overlap between the golimumab molecules and the lipid bilayer. As shown in **Fig. S18b**, we found a strong overlap between the bilayer and the golimumab bound to the ECD monomer engaged with the membrane (the minimum heavy atom distance measured 0.45 Å). This analysis provides a rationale to hypothesize that the steric obstruction mechanism, at least in part, could be responsible for reduced activity of biologics towards mTNF. Taken together with these experimental observations, our simulation results highlight the critical role of the membrane environment in modulating the conformational preferences and accessibility of mTNF to the receptor and therapeutic molecules.

We note that experimental validation of the key interaction mode of mTNF with the membrane identified in this work will be needed to further support its potential mechanistic relevance to TNF biology. Such experiments could be guided by our computational predictions on specific mutations that could potentially disrupt these membrane interactions (e.g., charge-neutralizing mutations of the key Arg residues in the ECD domain).

In summary, the presented work shows that the plasma membrane modulates the structural dynamics and energetic landscape of TNF (see **Fig. 8**). Our results offer a mechanistic hypothesis for the future design of therapeutics that selectively target sTNF or mTNF by exploiting the distinct conformational landscapes and membrane-dependent energetics of each form.

## SUPPORTING INFORMATION

Seventeen figures illustrating additional results.

## DATA AVAILABILITY

At the following URL, https://zenodo.org/uploads/22750222, we provide the starting and final conformations of the presented unbiased and meta-eABF simulations. We also provide the MD configuration files for the unbiased, meta-eABF, and implicit solvent MD simulations, alongside with the respective rst7 and parm7 files to reproduce the simulations.

## CODE AVAILABILITY

At the following URL, https://zenodo.org/uploads/22750222, we provide the analysis and run script files to reproduce the simulations and the data presented in this work.

## AUTHOR INFORMATION

## AUTHOR CONTRIBUTIONS

G.K. and F.T. designed research. G.K., F.T. and I.K performed experiments. G.K. analyzed data. G.K., F.T., and I.K. contributed analytic tools, interpreted the results, and wrote the manuscript. W.S. reviewed and edited the manuscript. All authors approved the final version of the manuscript.

## ACKNOWLEDGMENTS

We thank James O’Connell for useful discussions about TNF soluble and membrane forms, and Wonpil Im and his team for their guidance with using MolCube. This work was funded by Psivant Therapeutics.

## DECLARATION OF INTERESTS

I.K. and F.T. were employees of Psivant Therapeutics (now PsiThera) at the time this work was performed. W.S. is an employee of PsiThera. G.K. was a paid consultant for Psivant Therapeutics. PsiThera is developing modulators of tumor necrosis factor superfamily targets, including soluble TNFα. No patents or patent applications relate to the work described here. The study was designed, performed, and reported by the authors, whose respective roles are given in the Author Contributions section above.

## Notes

### Summary of Updates

Added simulation convergence analysis and improved manuscript.

