## Supplementary Material for "Interactions with lipid membrane modulate the conformational dynamics and energy landscape of Tumor Necrosis Factor"

### SUPPLEMENTARY FIGURES

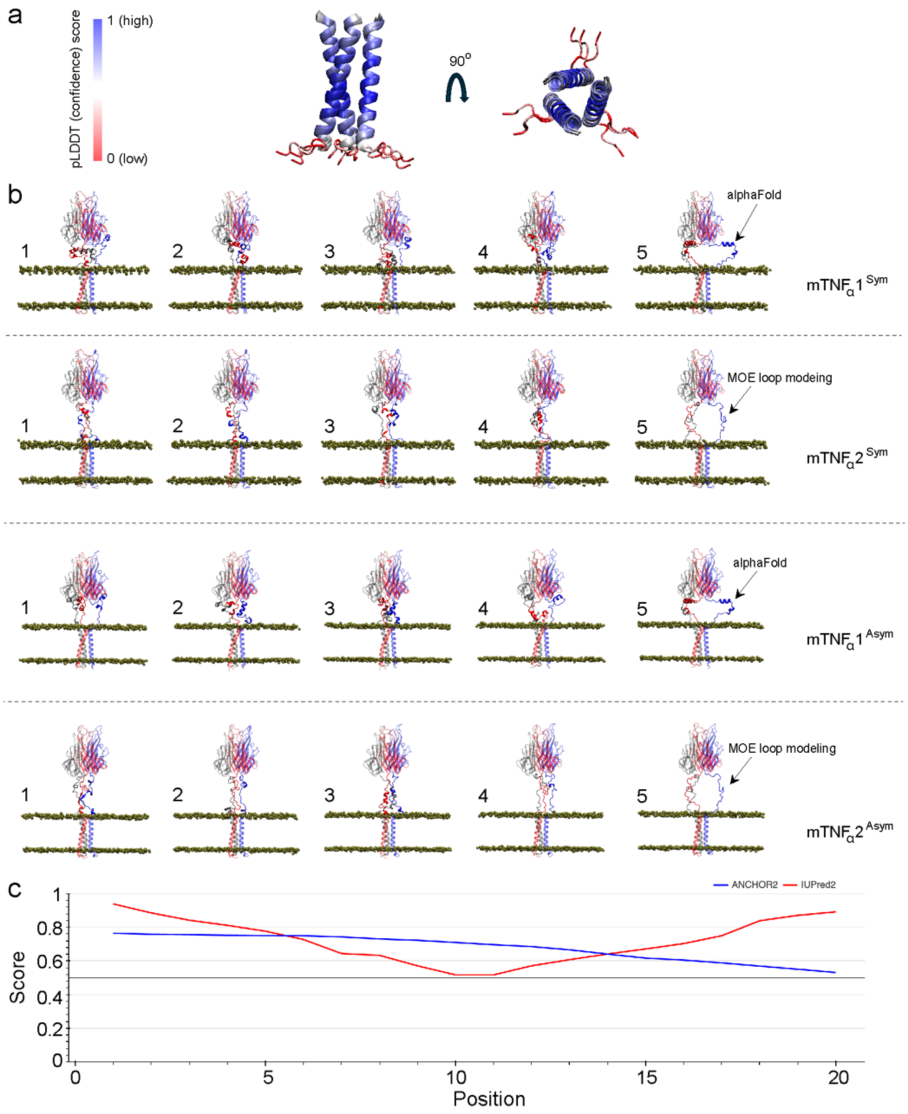


**Figure S1: Modeling of mTNF. (a)** Alphafold2 prediction of the TNF transmembrane domain. The pLDDT score indicates an overall high confidence that the transmembrane domain is trimeric. **(b)** Atomistic models of mTNF used as starting points for the unbiased MD simulations. The 5 models for mTNF1$^{Sym}$, mTNF2$^{Sym}$, mTNF1$^{Asym}$, and mTNF2$^{Asym}$ systems are shown separately in different rows. The replicates for a specific system are different in their structure of the linker loop connecting the ECD to the transmembrane bundle. The linker model predicted by AlphaFold are shown in replicate 5 for each system. The other 4 models are based on the clustering analysis of the implicit solvent MD simulation data of the AlphaFold-predicted loop and represent the centroid structures of the top 4 clusters (see Methods). **(c)** Disorder prediction of residues 1 to 20, performed through the web server https://iupred3.elte.hu. Scores above 0.5 indicate disordered residues. The two different colors indicate two different prediction models.

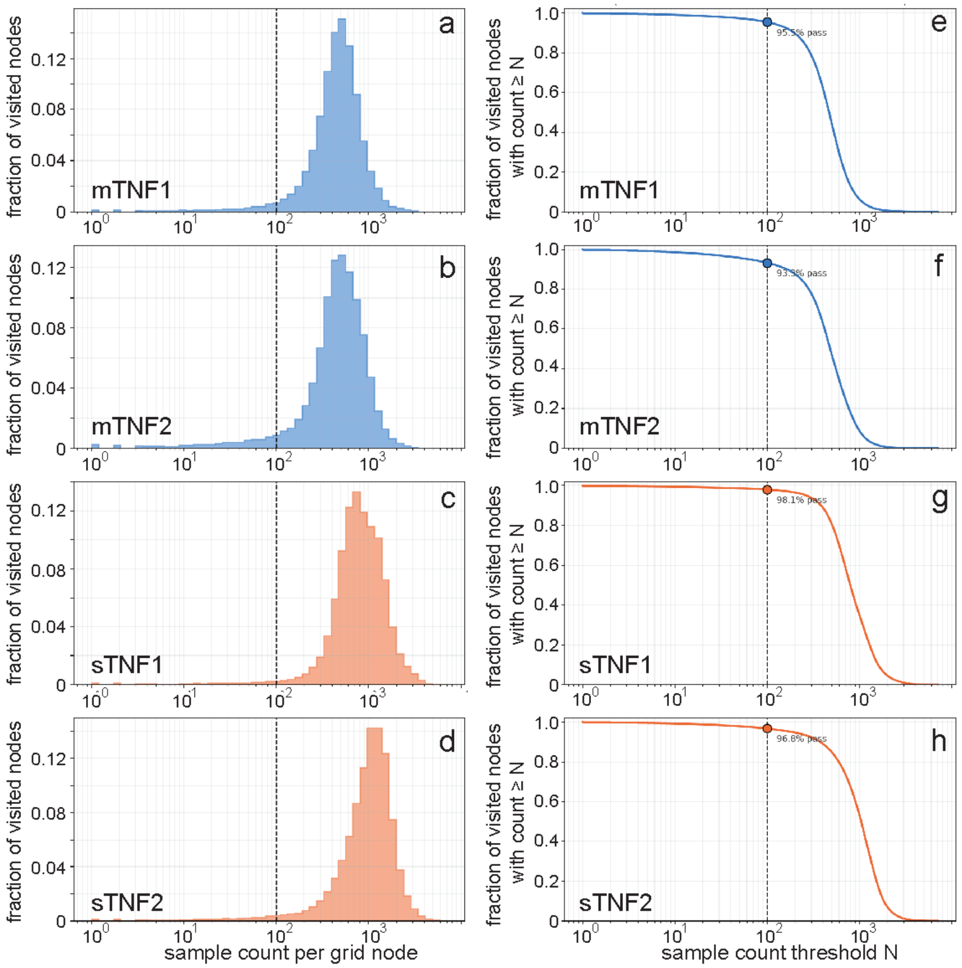


i

| STATE | MEAN NODE CZAR COUNTS | | | |
|---|---|---|---|---|
| | mTNF Rep1 | mTNF Rep2 | sTNF Rep1 | sTNF Rep2 |
| Symmetric | 1350 | 825 | 1670 | 691 |
| Asymmetric 1 | 508 | 682 | 1252 | 875 |
| Asymmetric 2 | 486 | 694 | 1105 | 672 |
| Asymmetric 3 | 439 | 916 | 2120 | 988 |

**Figure S2: Distribution and adequacy of per-bin CZAR sampling during meta-eABF**. **(a-d)** The distribution of per-node CZAR sample counts among grid nodes visited at least once, normalized to the fraction of visited nodes and shown on a log-scaled count axis for mTNF Rep1 (a), mTNF Rep2 (b), sTNF Rep1 (c), and sTNF Rep2 (d) systems. (**e-h**) The corresponding CZAR sampling-adequacy (survival) curves, giving the fraction of visited nodes retaining a sample count of at least N as a function of the threshold N. The filled circles in panels e-f mark the count ≥ 100 threshold used to mask all free-energy calculations in this work, with the retained fraction of visited nodes annotated at that point. The dashed vertical lines in panels a-f mark this same threshold. All four systems retain the large majority of their sampled domain at this threshold – 95.5% (mTNF Rep1), 93.3% (mTNF Rep2), 98.1% (sTNF Rep1), and 96.8% (sTNF2) of visited nodes – with median per-node counts of 468, 480, 792, and 1041 samples, respectively, confirming that the reported free energies rest on adequately sampled regions of CV space rather than a small number of sparsely visited outlier bins. (**i**) Mean CZAR sample counts for the nodes defining the regions of symmetric and 3 asymmetric states (see also Figure 7 table) for the 4 systems.

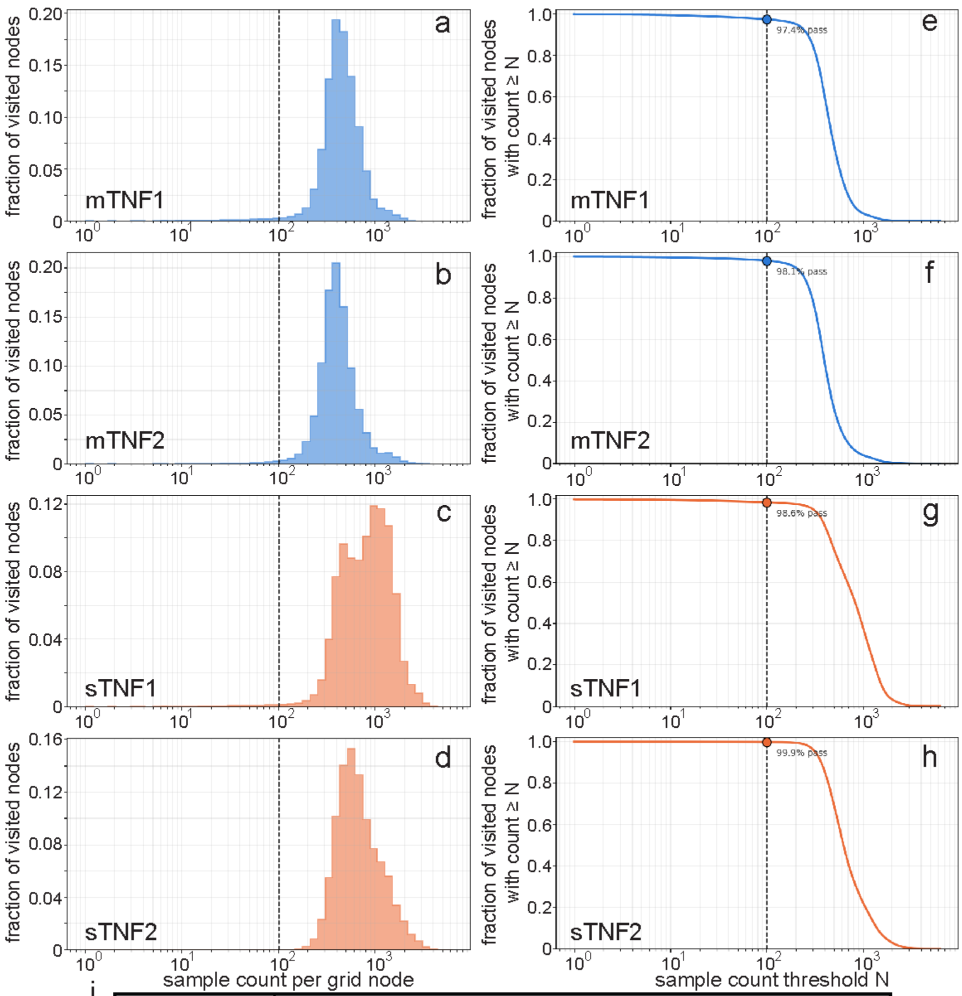


i

| STATE | MEAN NODE ABF COUNTS | | | |
|---|---|---|---|---|
| | mTNF Rep1 | mTNF Rep2 | sTNF Rep1 | sTNF Rep2 |
| Symmetric | 371 | 580 | 1016 | 450 |
| Asymmetric 1 | 347 | 359 | 961 | 958 |
| Asymmetric 2 | 487 | 397 | 904 | 562 |
| Asymmetric 3 | 511 | 361 | 1278 | 781 |

**Figure S3: Distribution and adequacy of per-bin ABF sampling during meta-eABF**. **(a-d)** The distribution of per-node ABF sample counts among grid nodes visited at least once, normalized to the fraction of visited nodes and shown on a log-scaled count axis for mTNF Rep1 (a), mTNF Rep2 (b), sTNF Rep1 (c), and sTNF Rep2 (d) systems. (**e-h**) The corresponding ABF sampling-adequacy (survival) curves, giving the fraction of visited nodes retaining a sample count of at least N as a function of the threshold N. The filled circles in panels e-f mark the count ≥ 100 threshold used to mask all free-energy calculations in this work, with the retained fraction of visited nodes annotated at that point. The dashed vertical lines in panels a-f mark this same threshold. All four systems retain the large majority of their sampled domain at this threshold – 97.4% (mTNF Rep1), 98.1% (mTNF Rep2), 98.6% (sTNF Rep1), and 99.9% (sTNF2) of visited nodes – with median per-node counts of 437, 395, 828, and 626 samples, respectively, confirming that the reported free energies rest on adequately sampled regions of CV space rather than a small number of sparsely visited outlier bins. (**i**) Mean ABF sample counts for the nodes defining the regions of symmetric and 3 asymmetric states (see also Figure 7 table) for the 4 systems.

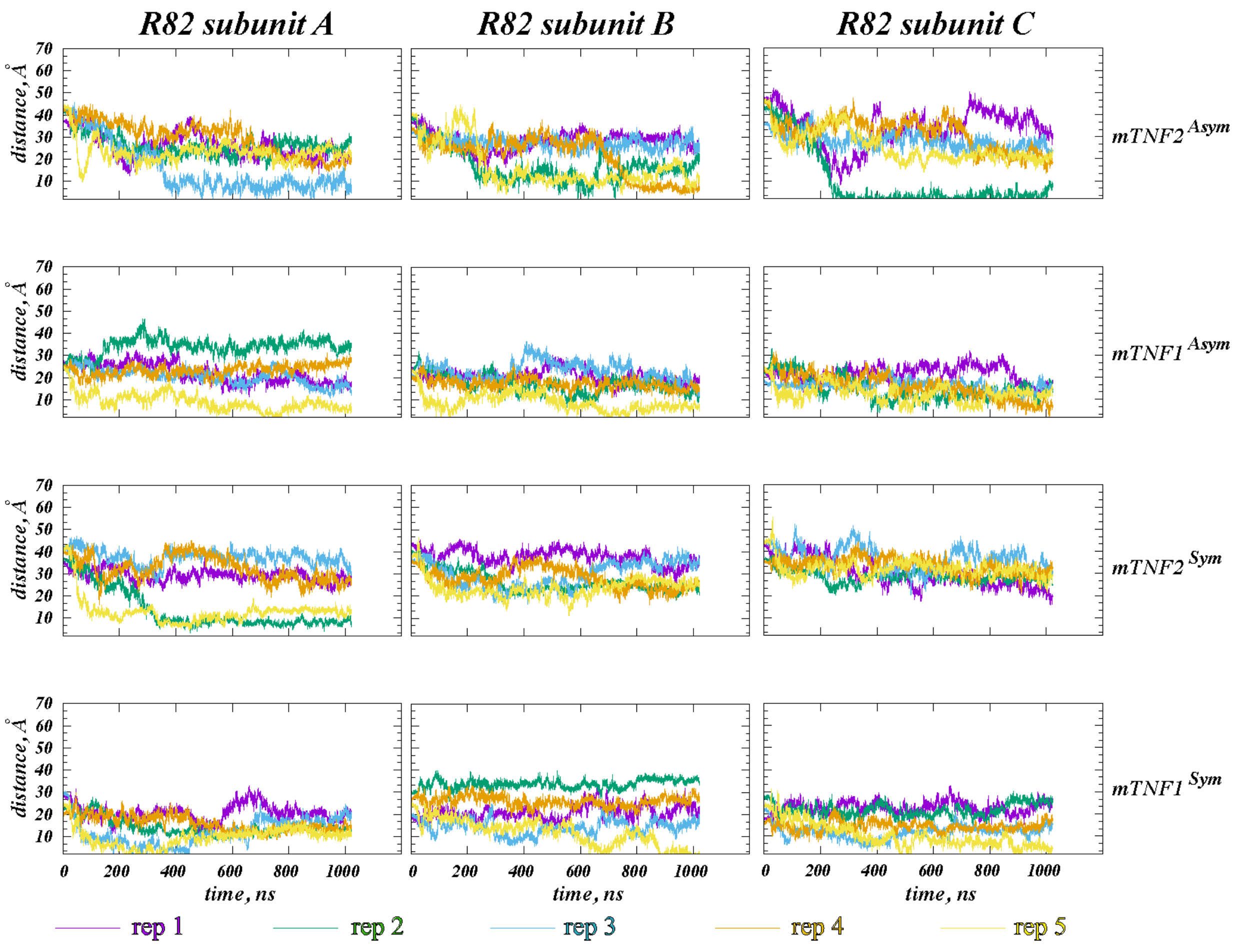


**Figure S4: Dynamics of membrane binding of residue R82 during the unbiased MD simulations.** Time evolution of the vertical (z-directional) distance between the $C_\alpha$ atom of residue R82 and the phosphorus atoms of the extracellular membrane leaflet. The data for separate subunits and for separate constructs are shown in different columns and rows, respectively. For each construct, the data for the five independent replicates are shown in different colors.

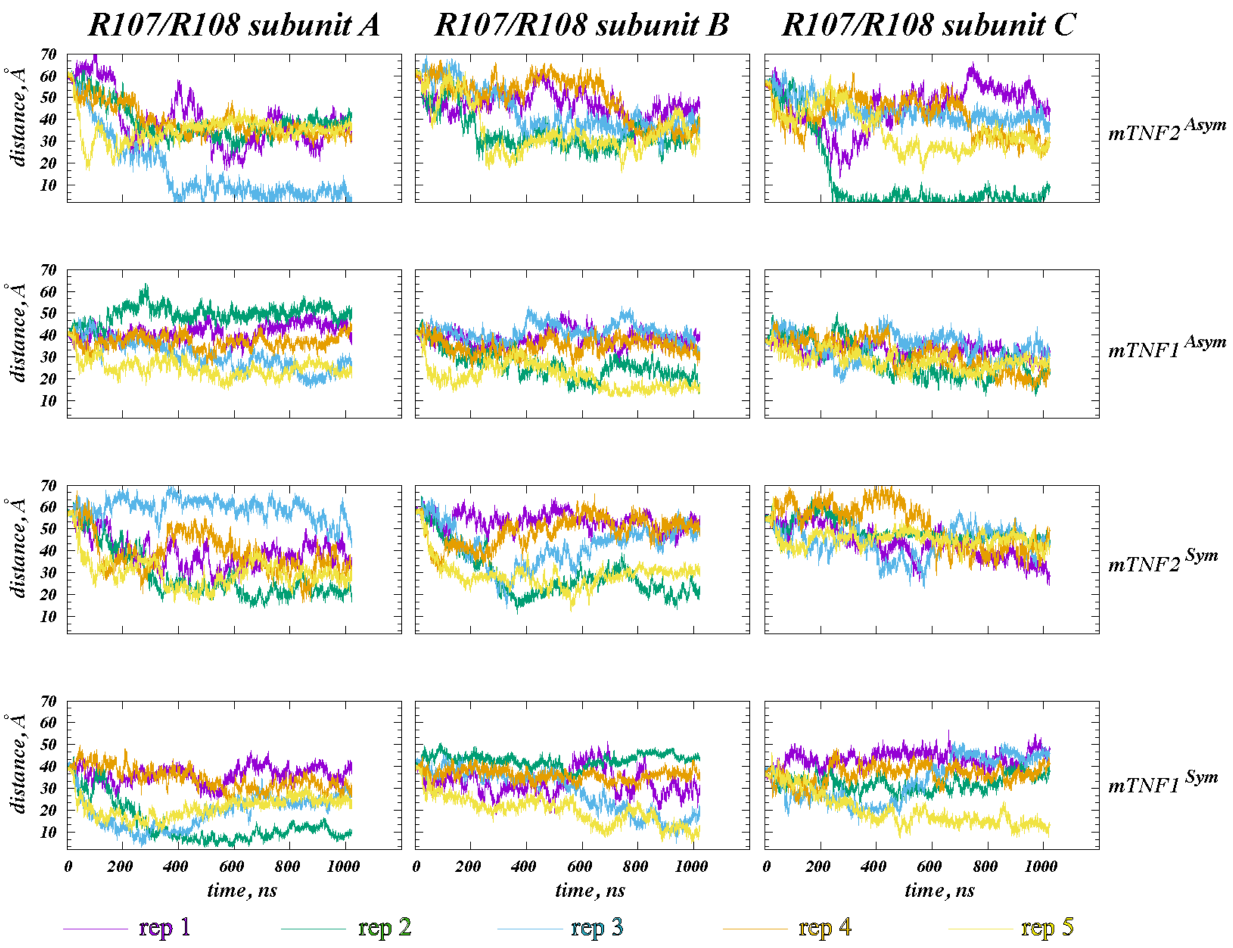


**Figure S5: Dynamics of membrane binding of residues R107/R108 during the unbiased MD simulations.** Time evolution of the vertical (z-directional) distance between the $C_\alpha$ atoms of residues R107/R108 and the phosphorus atoms of the extracellular membrane leaflet. The data for separate subunits and for separate constructs are shown in different columns and rows, respectively. For each construct, the data for the five independent replicates are shown in different colors.

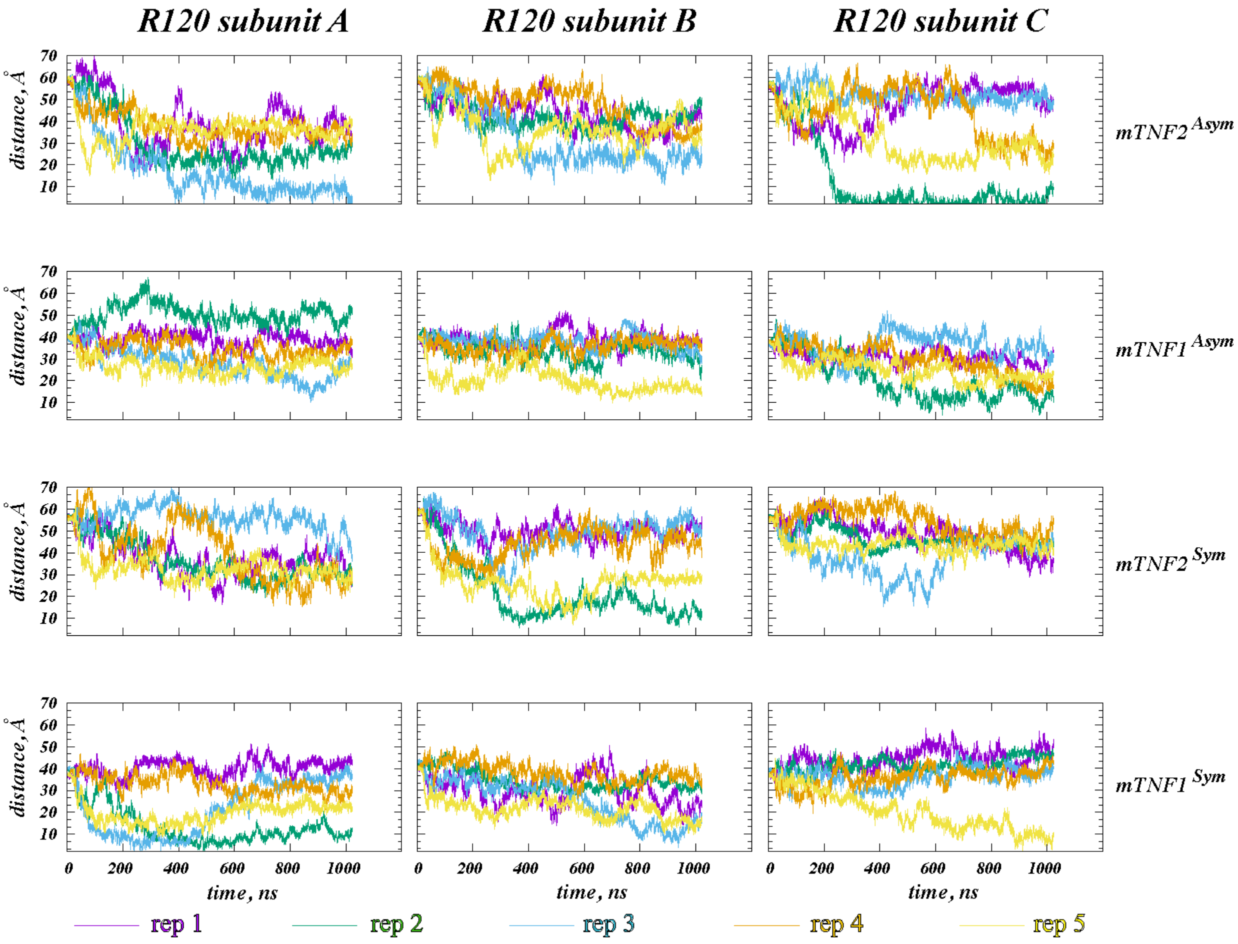


**Figure S6: Dynamics of membrane binding of residue R120 during the unbiased MD simulations.** Time evolution of the vertical (z-directional) distance between the $C_\alpha$ atom of residue R120 and the phosphorus atoms of the extracellular membrane leaflet. The data for separate subunits and for separate constructs are shown in different columns and rows, respectively. For each construct, the data for the five independent replicates are shown in different colors.

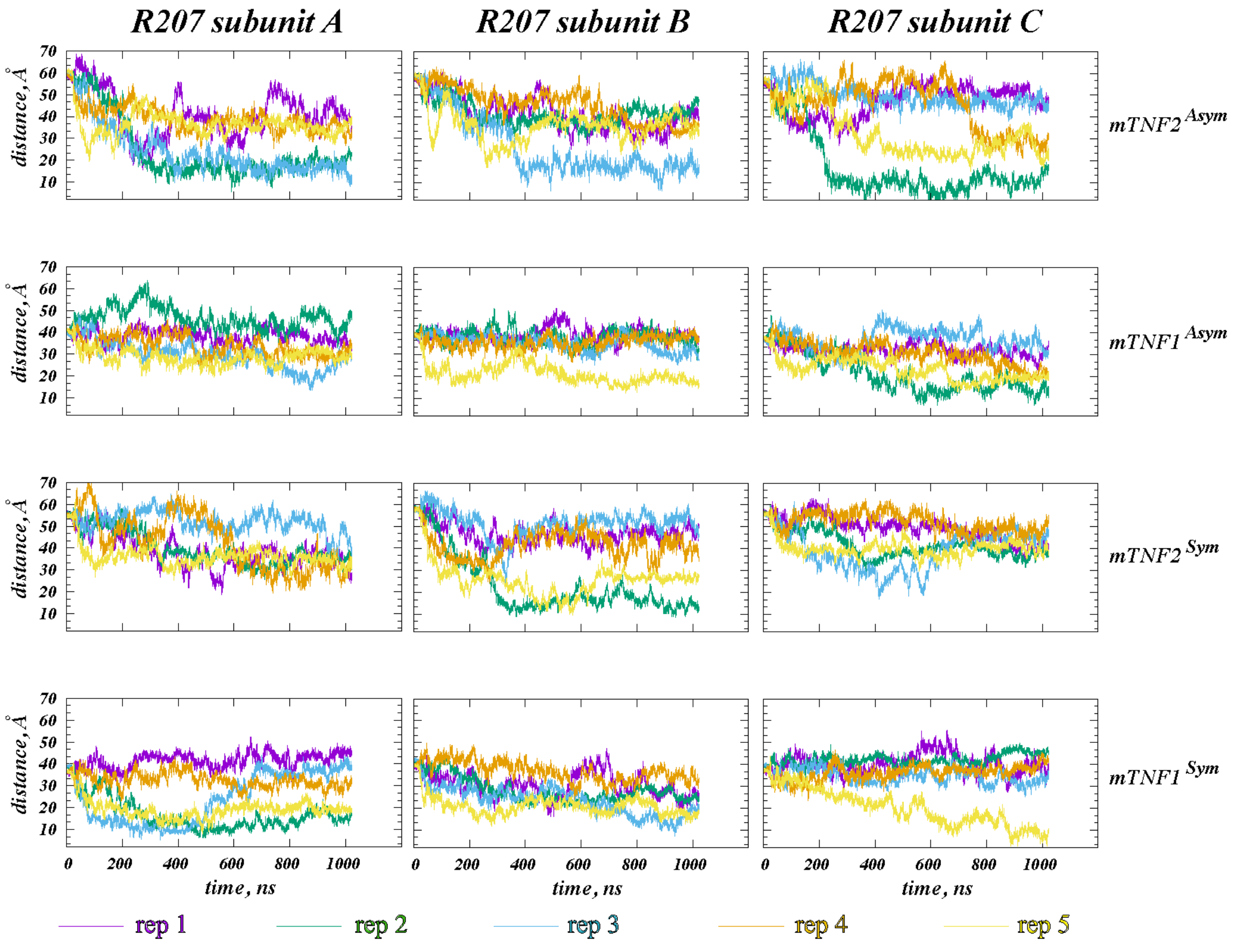


**Figure S7: Dynamics of membrane binding of residue R207 during the unbiased MD simulations.** Time evolution of the vertical (z-directional) distance between the $C_\alpha$ atom of residue R207 and the phosphorus atoms of the extracellular membrane leaflet. The data for separate subunits and for separate constructs are shown in different columns and rows, respectively. For each construct, the data for the five independent replicates are shown in different colors.

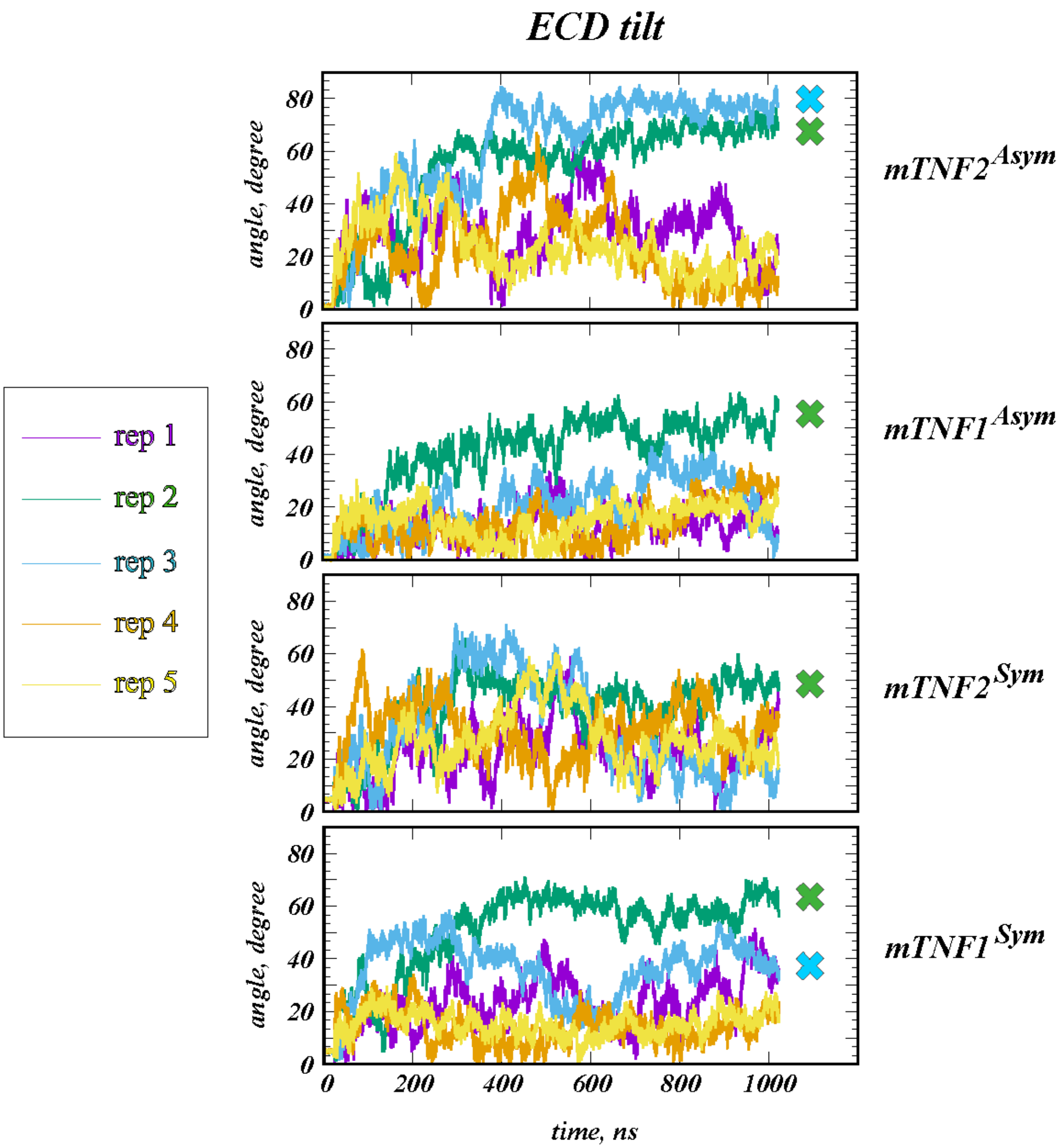


**Figure S8: Tilting of mTNF$_\alpha$ ECD during the unbiased MD simulations.** Time evolution of the tilt angle of the ECD with respect to the vertical axis. The data for separate constructs are shown in different panels. For each construct, the data for the five independent replicates are shown in different colors. The tilt angle was defined as the angle between the vertical axis and the vector connecting the center-of-masses of the C$_\alpha$ atoms of residues 233 and 176 from the 3 subunits. The simulations in which the Mode 1 membrane binding mode was observed is marked by "x" symbol (see also Table 1 of the main text).

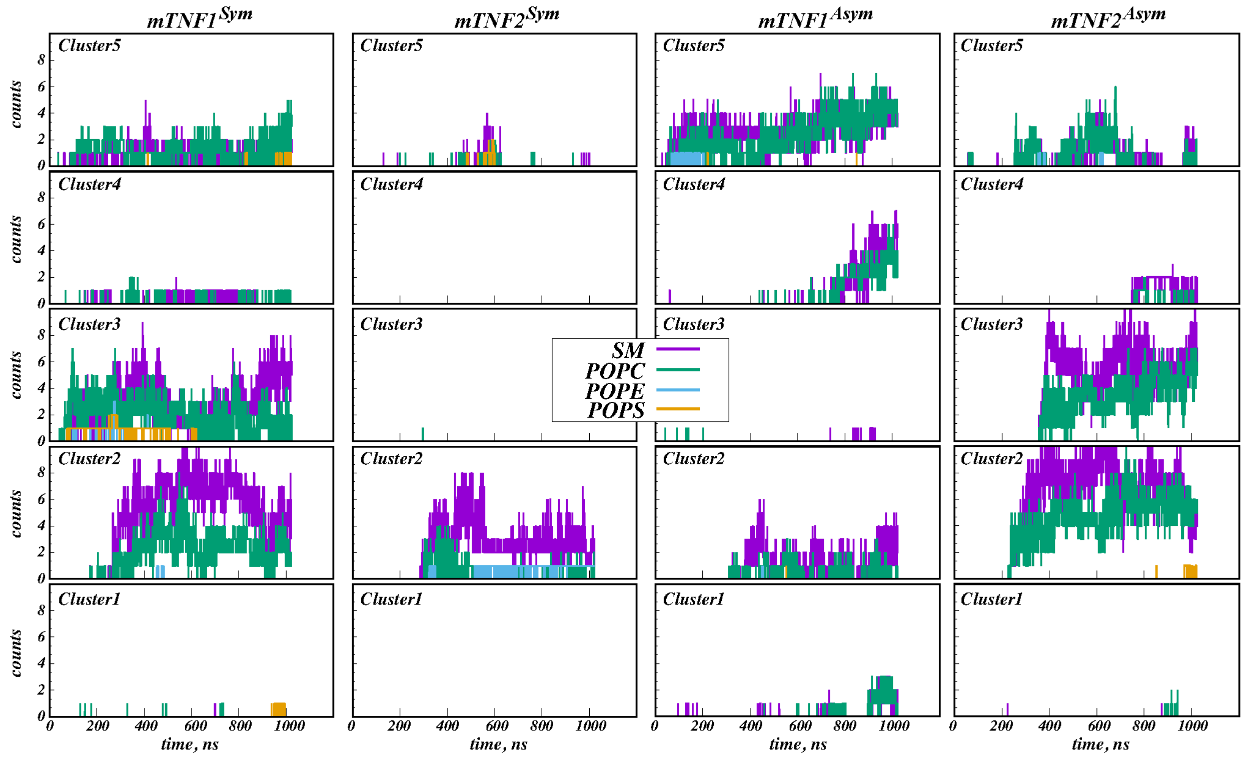


**Figure S9: EC domain of mTNF$_\alpha$ interacts mainly with POPC and SM lipids abundantly present on the EC membrane leaflet.** Time evolution in ensemble MD simulations of number of contacts between the EC domain of mTNF$_\alpha$ and individual lipid species (SM, POPC, POPE, and POPS). A contact was assumed if a distance between any of the atoms on residues R82, R107, R108, R120, and R207 was within 4Å of any of the lipid atom.

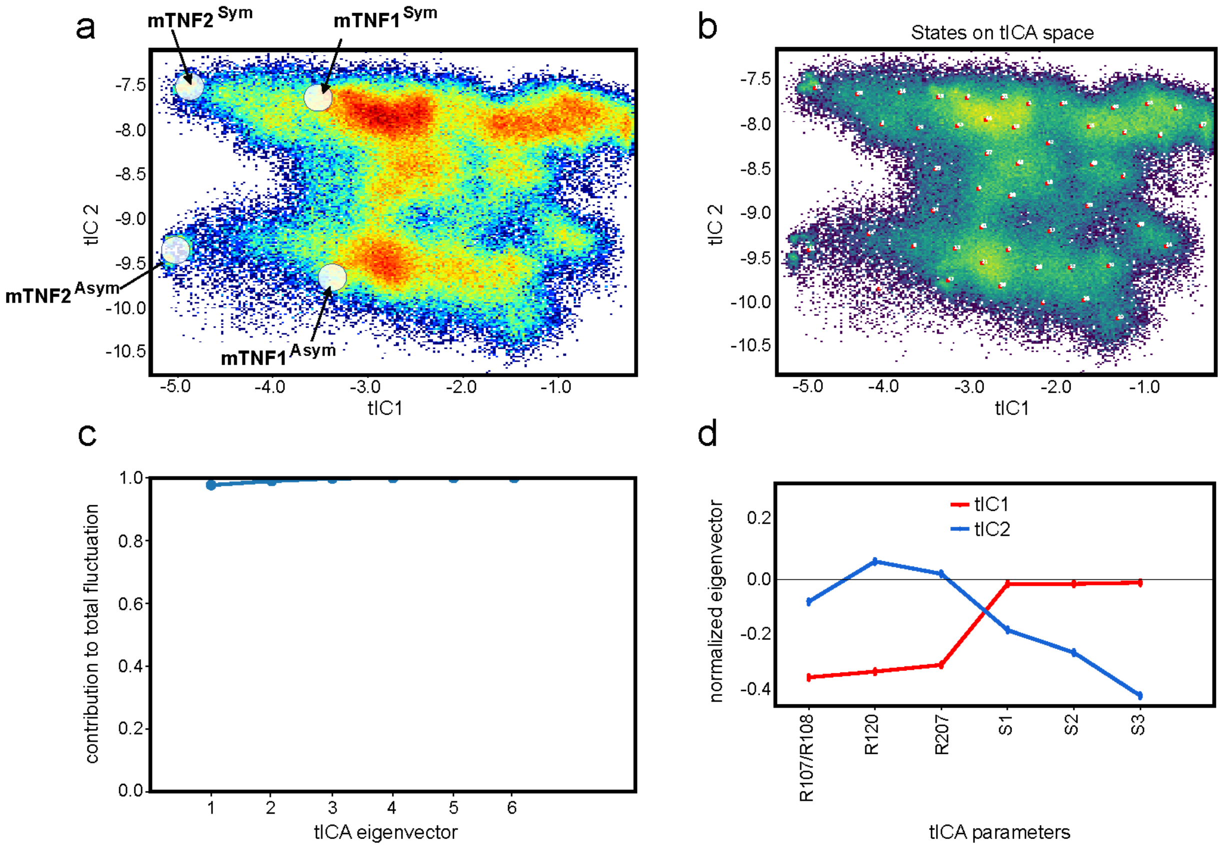


**Figure S10: tICA dimensionality reduction of the unbiased MD simulations of mTNF. (a)** Projection of all the trajectory frames from the unbiased MD simulations onto the 2D space spanned by the first two tIC vectors (from Fig. 4a of the main text). The projections of the 4 initial structural models on the tICA space are indicated. **(b)** The centers of the most populated states on the tICA landscape (indicated by the numbered circles) obtained as a result of the tICA space clustering (see Methods). **(c)** Contributions of the tICA eigenvectors to the total fluctuations in the system. **(d)** Weights of the structural features (CVs) to the first two tIC vectors, tIC1 (in red) and tIC2 (in blue).

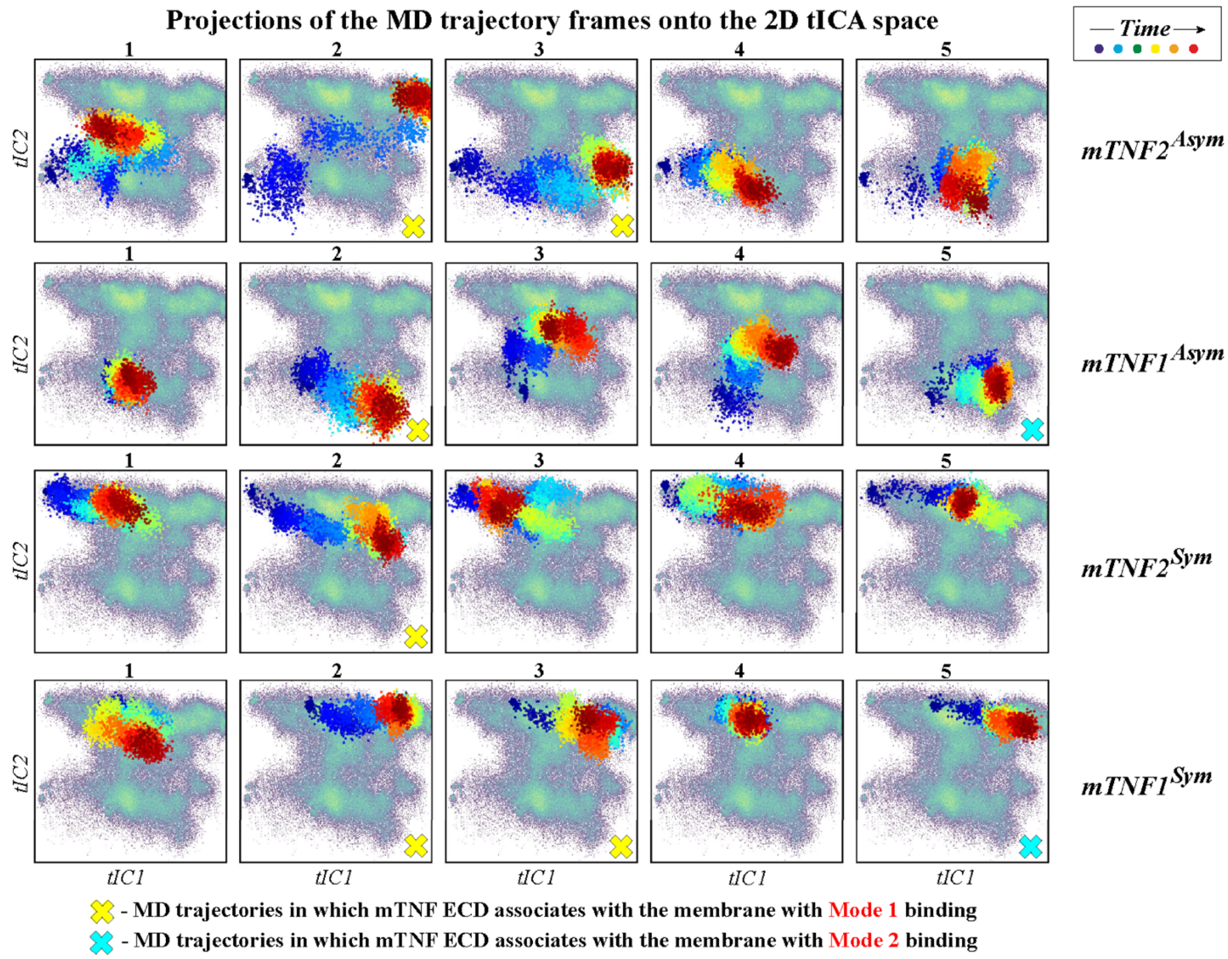


**Figure S11: Projections of the individual MD trajectories onto the tICA landscape.** Projection (colored dots) of each MD trajectory from all the replicates from the 4 sets of simulations on the 2D tICA landscape (in pale colors) from Fig. 4. The colors of the dots indicate the timeframes in the evolution of the trajectory: darker colors (blue, cyan) represent the initial stages of the simulation, lighter colors (yellow, green) correspond to the middle part of the trajectory, and reds show the last third of the trajectory. The replicates in which the ECD associated with the membrane with Mode 1 and Mode 2 binding are marked by yellow and cyan "x" symbols.

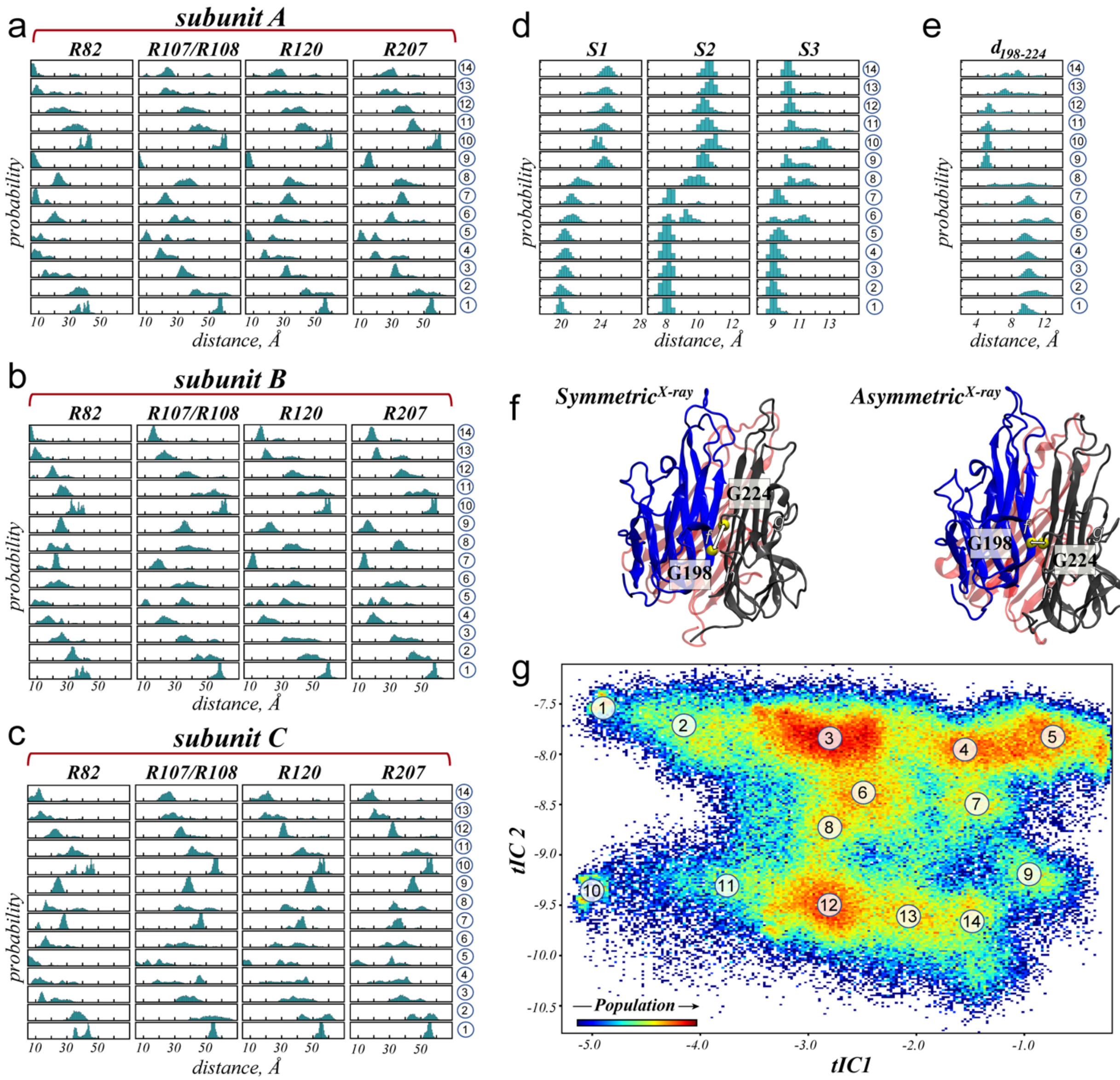

**Figure S12: Structural characteristics of the tICA landscape. (a-c)** The histograms quantifying membrane proximity of the selected basic residues on subunit A (a), subunit B (b), and subunit C (c) of the ECD. The plots show probabilities in selected states on the tICA space (shown in panel g) of vertical (z-directional) distance of the $C_{\alpha}$ atoms of residues R82, R107/108, R120, and R207 to the lipid phosphorus atoms in the extracellular leaflet. **(d)** Histograms of S1, S2, and S3 distances in the selected states of the tICA space (shown in panel g). **(e)** Histograms of the $C_{\alpha}$-$C_{\alpha}$ distance $d_{198-224}$ between residues G198 and G224 in the selected states of the tICA space (shown in panel g). **(f)** Visual representation of the Symmetric$^{X\text{-ray}}$ and Asymmetric$^{X\text{-ray}}$ structures showing different positioning of G198 and G224 residues in the two states. The *f* strand of the β-sheet motif in subunit A and the *g* and *h* strands of the β-sheet motif in subunit C are indicated. **(g)** Projection of all the trajectory frames from the unbiased MD simulations onto the 2D space spanned by the first two tIC vectors. The color map identifies the populations distribution, and the locations of selected states on the 2D space are indicated by numbers from 1 through 14.

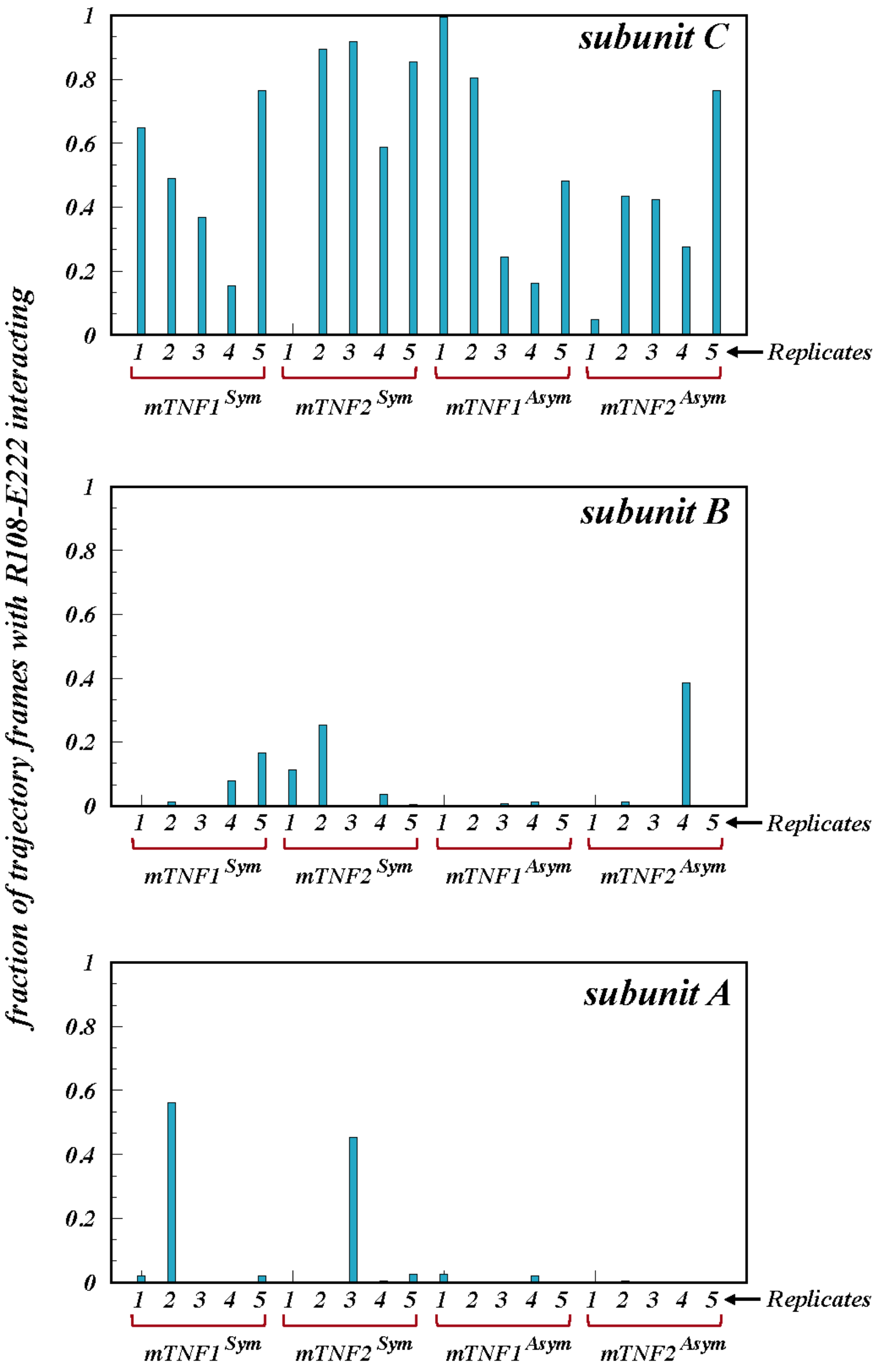


**Figure S13: Interactions between R108 and E222 during unbiased MD simulations.** Fraction of trajectory frames with R108-E222 interacting shown for each replicate and from all the 4 sets of simulations. The data for subunits A, B, and C are shown in bottom, middle, and top panels, respectively.

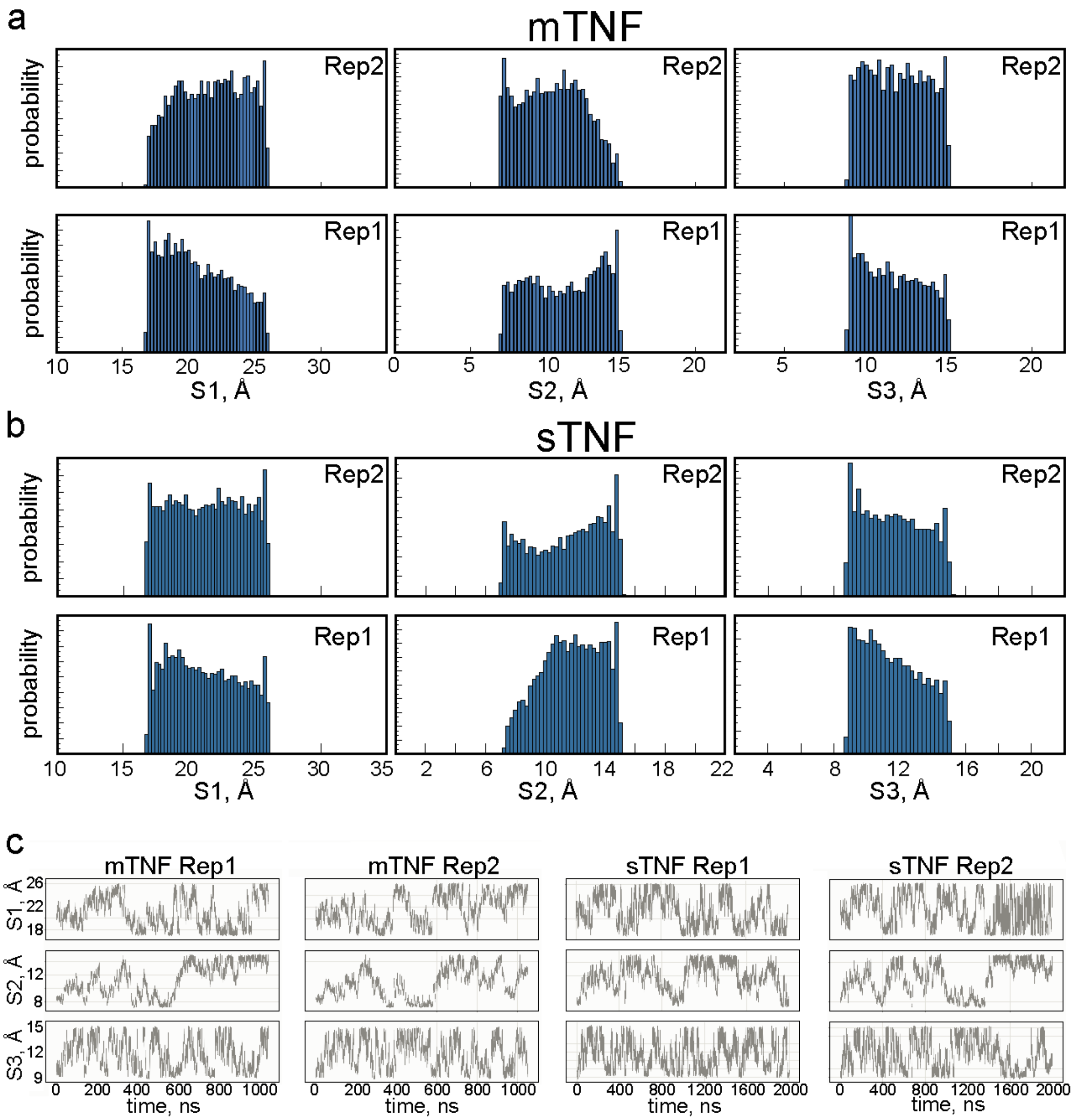


**Figure S14: Sampling of the CVs in the meta-eABF MD simulations. (a-b)** Histograms of the S1, S2, and S3 variables in the meta-eABF simulations of mTNF (a) and sTNF (b) systems. The data for 2 independent replicates per construct are shown in separate rows. (**c**) Time-traces of the S1, S2, and S3 variables in the meta-eABF simulations of mTNF and sTNF systems. The data for 2 independent replicates per construct are shown in separate columns.

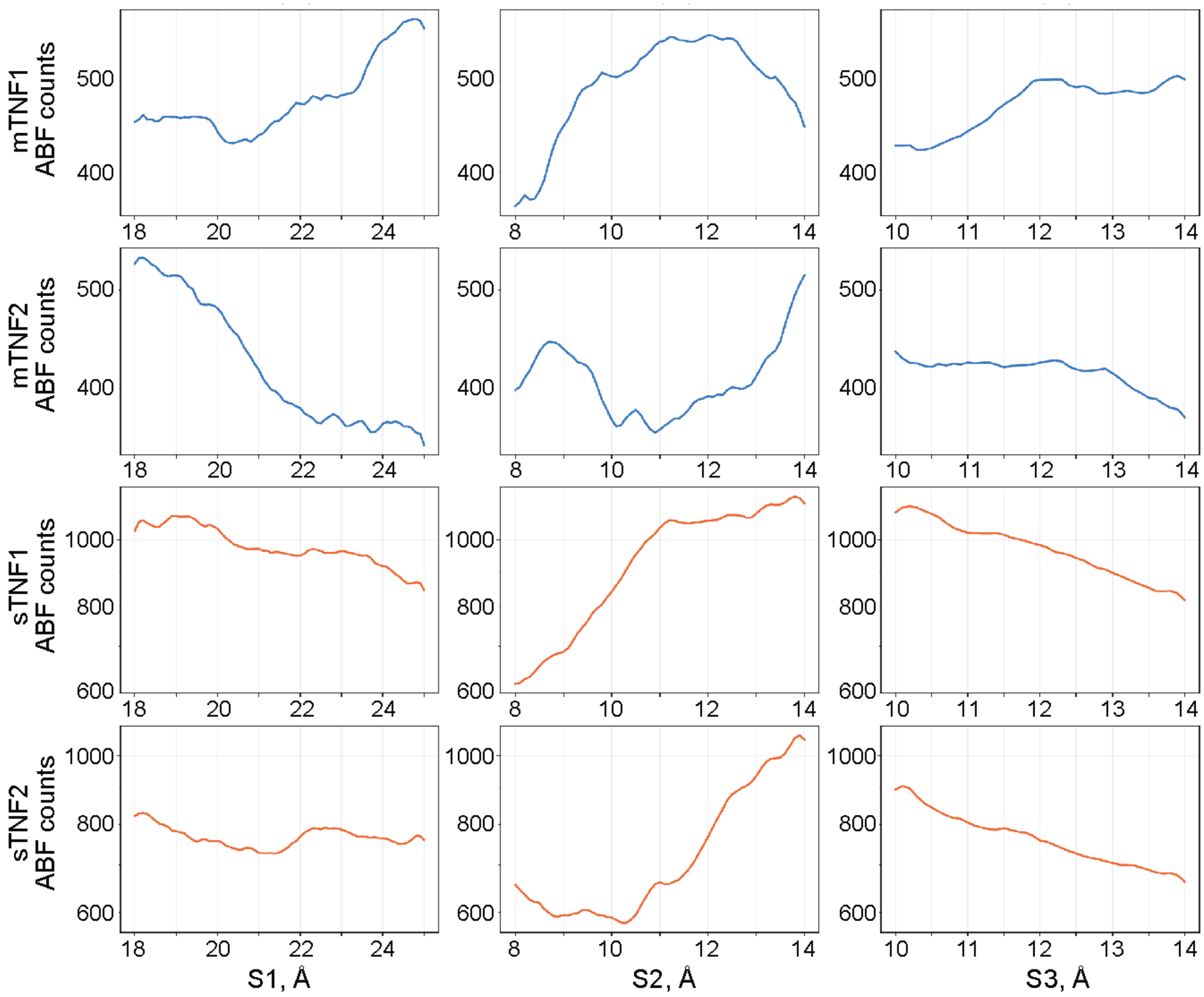


**Figure S15: ABF sample-count flatness as a diffusive-sampling check.** For each system, the ABF module's raw grid sample count averaged over the two collective variables not shown, restricted to grid nodes passing the count ≥ 100 adequacy threshold used throughout this work, and plotted (log scale) against the third collective variable (S1, S2, S3, left to right). Rows show mTNF Rep1, mTNF Rep2, sTNF Rep1, and sTNF Rep2 (top to bottom). The profiles are relatively flat (within roughly a factor of 1.5-2.0) in all four systems and for all three CVs, indicative of reasonable convergence.

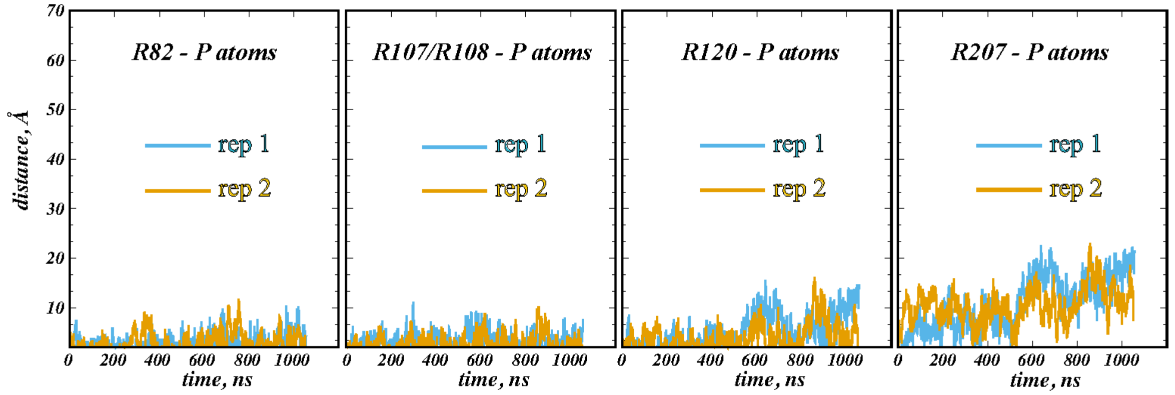


**Figure S16: ECD-membrane interactions during meta-eABF simulations.** Time evolution of the vertical (z-directional) distance between the $C_\alpha$ atoms of residues R82, R107/R08, R120, and R207 and the lipid phosphorus atoms in the extracellular membrane leaflet in the meta-eABF simulations of mTNF.

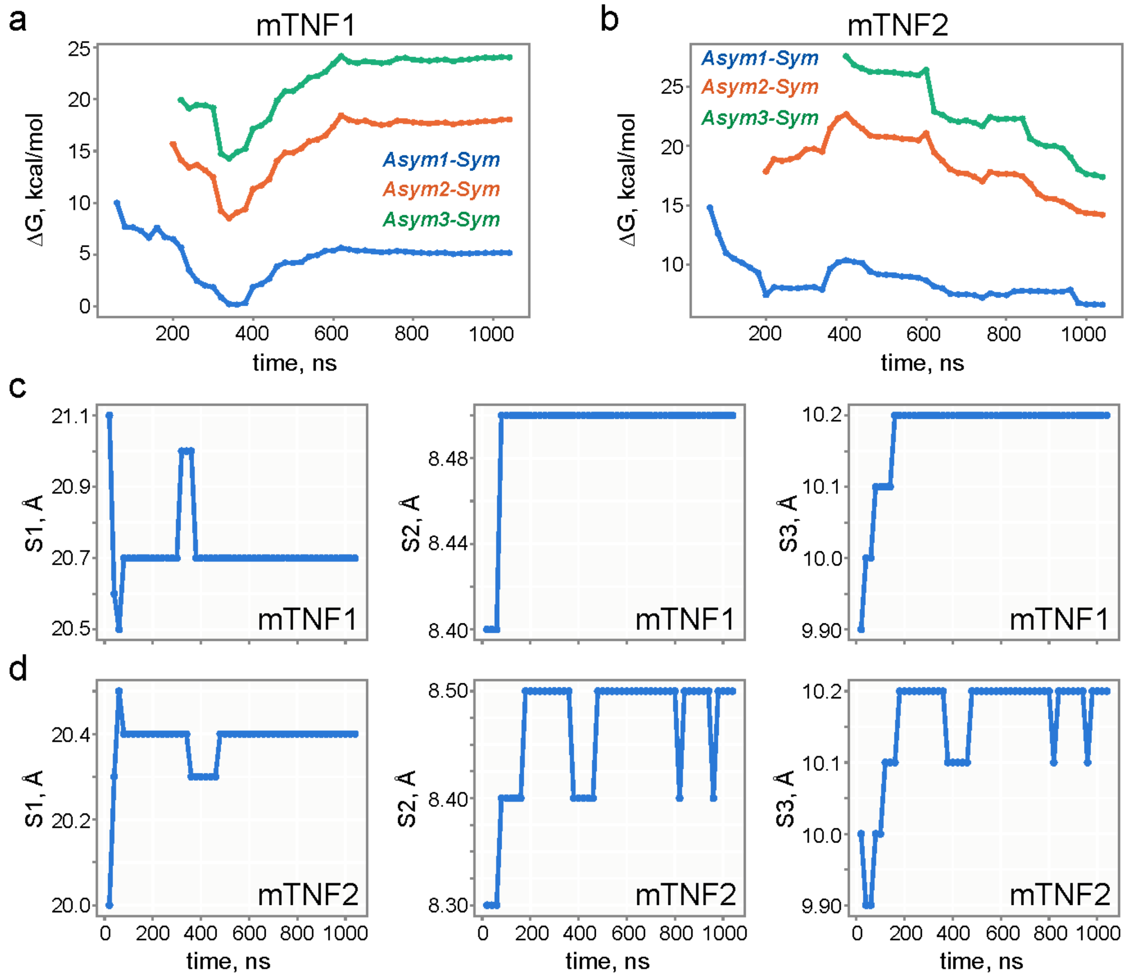


**Figure S17: Convergence of the meta-eABF free energy estimates. (a-b)** Cumulative estimates of the free energy differences between the main Asymmetric states 1, 2 and 3 (Asym1-3) and the Symmetric (Sym) state, $\Delta G_{Asym1-Sym}$ (blue), $\Delta G_{Asym2-Sym}$ (orange), and $\Delta G_{Asym3-Sym}$ (green), as a function of trajectory time for mTNF1(a) and mTNF2 (b) systems. The profiles start at the time points when the respective basins accumulate ≥ 1 sampled bin passing the count threshold (see Methods). All three estimates for both systems stabilize to within the final plateau value for the remainder of the trajectory, indicating convergence of the three quantities. (**c-d**) The collective variable coordinates (S1, S2, S3 from left to right) of the single lowest-free-energy grid node identified from the cumulative trajectory up to the specific time, plotted against the simulation time for mTNF1 (c) and mTNF2 (d) systems. The identified minimum settles onto a fixed grid node and remains there for the remainder of the simulation in both systems, indicating convergence of the global free-energy basin's location.

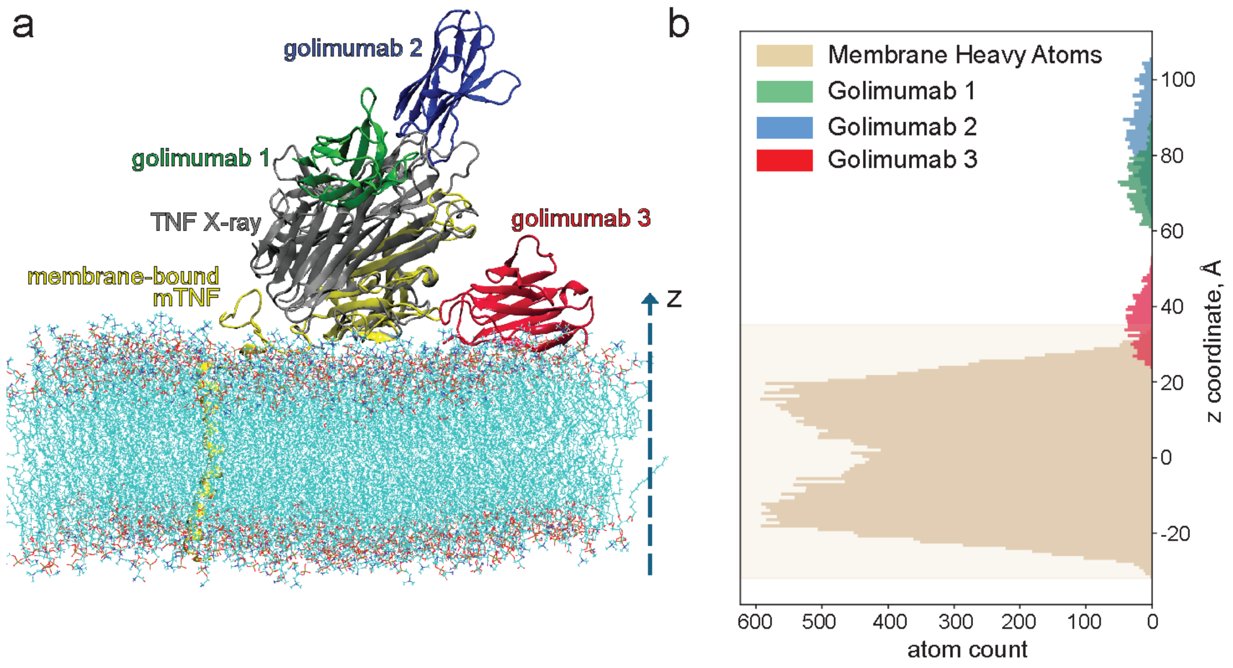


**Figure S18: Evaluation of potential steric clashes between TNF-bound biologics and the membrane when mTNF engages in Mode 1 binding with the lipid membrane. (a)** A snapshot showing structural alignment of the X-ray model of sTNF in complex with golimumab biologics (PDBID: 5YOY) and the membrane-bound mTNF from MD simulations. The latter represents Mode 1 binding. The 3 golimumab molecules are colored in blue, green, and red. The sTNF from the X-ray model is in grey, and the membrane-bound mTNF chain is shown in yellow (the two remaining chains are omitted for clarity). The membrane is depicted as lines. (**b**) Plot of the number of membrane heavy atoms (gold), and of the three golimumab molecule atoms (green, blue, and red) found at specified z coordinate (membrane-normal axis), showing an overlap between one of the golimumab and the membrane. The minimum distance between the overlapping golimumab molecule (in red) and the membrane is 0.45 Å, suggesting evere steric clashes.
